# Bronchoalveolar Lavage Single-Cell Transcriptomics Identifies Immune Cells Driving COVID-19 Severity in Patients

**DOI:** 10.1101/2024.08.22.609137

**Authors:** Clinton Njinju Asaba, Razieh Bitazar, Patrick Labonté, Terence Ndonyi Bukong

**Author notes:** Corresponding author: Terence Ndonyi Bukong.

## Abstract

The continuous threats posed by Severe Acute Respiratory Syndrome Coronavirus 2 (SARS-CoV-2), the virus that causes COVID-19, including the emergence of potentially more infectious and deadly variants, necessitate ongoing studies to uncover novel and detailed mechanisms driving disease severity. Using single-cell transcriptomics, we conducted a secondary data analysis of bronchoalveolar lavage fluid (BALF) from COVID-19 patients of varying severities and healthy controls to comprehensively examine immune responses. We observed significant immune cell alterations correlating with disease severity. In severe cases, macrophages showed upregulation of pro-inflammatory genes TNFα and IL1β, contributing to severe inflammation and tissue damage. Neutrophils exhibited increased activation, marked by S100A8, CXCL8, and IL1β expression, with extended viability and reduced phagocytosis. Genes such as MCL1 and HIF1α supported extended viability, while MSR1 and MRC1 indicated reduced phagocytosis. Enhanced formation of neutrophil extracellular traps (NETs) and reduced clearance, indicated by NET-associated markers, were linked to thrombo-inflammation and organ damage. Both macrophages and neutrophils in severe cases showed impaired efferocytosis, indicated by decreased expression of MSR1 and TREM2 in macrophages and downregulation of FCGR3B in neutrophils, leading to the accumulation of apoptotic cells and exacerbating inflammation. Severe cases were characterized by M1 macrophages with high TNFα and IL1β, while milder cases had M2 macrophages with elevated PPARγ. Low-density neutrophils (LDNs) increased significantly in severe cases, showing higher CXCR4 and CD274 and lower FCGR3B compared to high-density neutrophils (HDNs). NK and T cells in severe cases demonstrated altered receptor and gene expression, with increased activation markers IFNγ and ISG15, suggesting a paradoxical state of activation and exhaustion. This imbalance suggests a potential mechanism for immune dysregulation and ineffective antiviral responses in severe COVID-19. This analysis highlights the critical role of dysregulated neutrophil, macrophage, NK, and T cell responses in severe COVID-19, identifying potential therapeutic targets and providing novel insights into the disease.

**Author Summary:** Severe Acute Respiratory Syndrome Coronavirus 2 (SARS-CoV-2), the causative agent of COVID-19, poses continuous health threats due to emerging, potentially more infectious and deadly variants. We used single-cell gene analysis from lung fluid samples, known as bronchoalveolar lavage (BAL), from COVID-19 patients to understand why some cases become more severe than others. In severe cases, immune cells called macrophages and neutrophils showed higher levels of genes that trigger inflammation and cause damage to the body. These cells were more active and lived longer but were less capable of clearing away dead cells and debris, leading to prolonged inflammation. Severe cases also had more neutrophils that were less effective in fighting infections. Another type of immune cell, NK and T cells, showed changes indicating an ineffective response to the virus, with signals that were not properly coordinated to fight the infection. This imbalance in the immune response can lead to severe inflammation and organ damage. Our findings highlight potential targets for treatments to help manage severe COVID-19 and improve patient outcomes. Understanding these immune cell behaviors through single-cell gene analysis could guide the development of new therapies and improve strategies for treating severe COVID-19.

## Introduction

The COVID-19 pandemic, caused by the severe acute respiratory syndrome coronavirus 2 (SARS-CoV-2), has significantly impacted global health. As of July 14, 2024, over 775 million cases have been confirmed, with more than 7 million deaths reported[1]. This crisis has placed unprecedented strain on healthcare systems while also driving remarkable scientific collaboration, particularly in immunology. This pandemic underscores the urgent need to improve our understanding of viral pathogenesis and the immune system’s role in both driving and combating infectious diseases.

A key revelation from this pandemic is the central role played by the immune system in determining the severity and outcomes of SARS-CoV-2 infection[2, 3]. Despite significant advances, the complexities of the innate immune response, especially in regulating inflammation and tissue damage during SARS-CoV-2 infection, are not fully understood. Critical areas of focus include the roles of type 1 and type 2 innate lymphoid cells, NK cells, T cell regulation, and the production of type 1 interferons (Interferon alpha and beta) and type 2 interferon (Interferon gamma)[4]. Neutrophils and macrophages, key components of the innate immune system, are crucial in the early stages of infection and significantly influence disease progression and severity[4, 5].

Neutrophils, the most abundant white blood cells, act as frontline defenders against pathogens through mechanisms such as phagocytosis and the release of neutrophil extracellular traps (NETs)[6]. While these processes are vital for controlling pathogens, excessive neutrophil activation and NET formation can cause significant tissue damage, contributing to the severe inflammatory responses seen in critical COVID-19 cases[7–9]. This phenomenon, often referred to as a “cytokine storm,” is a hallmark of severe SARS-CoV-2 infection and illustrates the double-edged nature of the immune response[7, 10]. Macrophages, known for their versatility, exhibit a range of activation states from the pro-inflammatory M1 phenotype, which is involved in pathogen killing and promoting inflammation, to the anti-inflammatory M2 phenotype, which aids in tissue repair and the resolution of inflammation[11–13]. Maintaining a balance between these phenotypes is crucial; an imbalance skewed towards the M1 phenotype can lead to unchecked inflammation and significant tissue damage, which are characteristic of severe COVID-19[14]. Similarly, natural killer (NK) cells and their interactions with macrophages and T cells are vital in modulating immune responses, impacting viral clearance and tissue damage[15–18]. The production of type 1 and type 2 interferons by these cells plays a crucial role in antiviral defense and immune response regulation[19, 20]. Understanding this balance is essential for developing therapeutic strategies aimed at modulating the immune response to achieve better clinical outcomes.

Research indicates that alterations in receptor and interferon regulatory genes affect the activity of type 1 and type 2 innate lymphoid cells, NK cells, and T cells, which in turn influence neutrophil and macrophage functions and correlate with disease progression[21–27]. For example, the presence of low-density neutrophils with enhanced activation markers in severe cases points to a hyperactive state that likely exacerbates disease progression[28, 29]. Moreover, elevated activation markers in M1 macrophages indicate a robust inflammatory response, presenting potential therapeutic targets for modulating immune responses in COVID-19[30, 31]. The regulation of T cell responses, particularly the balance between effector and regulatory T cells, is also crucial in determining the outcome of SARS-CoV-2 infection[32, 33]. These insights not only enhance our understanding of SARS-CoV-2 pathogenesis but also underscore the urgent need for targeted therapeutic strategies to mitigate severe inflammatory responses and improve patient outcomes.

Our novel study demonstrates the transformative potential of single-cell transcriptomics in providing unprecedented resolution into complex immune interactions[34–36]. By offering a comprehensive framework for future research and therapeutic development, our work underscores the critical importance of advanced transcriptomic technologies in understanding and combating infectious diseases. This research paves the way for innovative therapeutic approaches that can be applied not only to COVID-19 but also to other viral infections, highlighting the broader relevance and impact of our findings in the field of host pathogen interactions.

## Material and Methods

### Ethics Statement

In this research, we conducted a secondary analysis using two publicly available datasets obtained from the Gene Expression Omnibus (GEO), a functional genomics data repository. As the participant data had already been anonymized and made publicly accessible, institutional review board (IRB) and ethical approval were not required. GEO supports MIAME-compliant data submissions, ensuring high standards for data quality and reporting. Since the datasets were fully anonymized, there were no additional ethical concerns regarding participant confidentiality or consent. Our approach aligns with established ethical guidelines, recognizing the minimal risk involved in analyzing de-identified public data, and complies with all relevant ethical standards while contributing valuable insights to the scientific community.

### Study Data

We acquired single-cell RNA sequencing (ScRNA-seq) data from bronchoalveolar lavage fluids (BALFs) of healthy individuals, moderately ill patients, and severe COVID-19 patients. These datasets were obtained from the Gene Expression Omnibus (GEO) database, accessed and downloaded between May 10^th^ and June 10^th^ 2024, which is owned and operated by the National Center for Biotechnology Information (NCBI), a part of the National Library of Medicine (NLM) at the National Institutes of Health (NIH) in the United States. The GEO database is a public archive and resource for gene expression data. Specifically, we used datasets with accession codes GSE151928 for healthy individuals, GSE145926 for moderately ill patients, and GSE157344 for severe COVID-19 patients. Our study included ScRNA-seq data from a total of 6 healthy controls, 3 moderately ill patients, and 21 severe COVID-19 patients. The GEO codes for healthy control individuals included: GSM4593888, GSM4593889, GSM4593890, GSM4593891, GSM4593892, GSM4593893. For moderate COVID-19 individuals the sample GEO codes for individuals used included: GSM4339769, GSM4339770, GSM4339772. For severe alive patients sample GEO codes included: GSM4762146, GSM4762141, GSM4762142,GSM4762143,GSM4762144,GSM4762155,GSM4762156, GSM4762157, GSM4762148, GSM4762159, GSM4762160, GSM4762139 and GSM4762151. For severe dead individuals samples analysed had the following GEO codes: GSM4762140, GSM4762153, GSM4762147, GSM4762149, GSM4762152, GSM4762158, GSM4762145, GSM4762150. A comprehensive summary of the basic biological and clinical information of the individuals and patients analyzed in this study can be found in **Table 1**. The data is fully anonymized, and we never had access to any identifiable information about the participants, either during or after data collection.

**Table 1.**
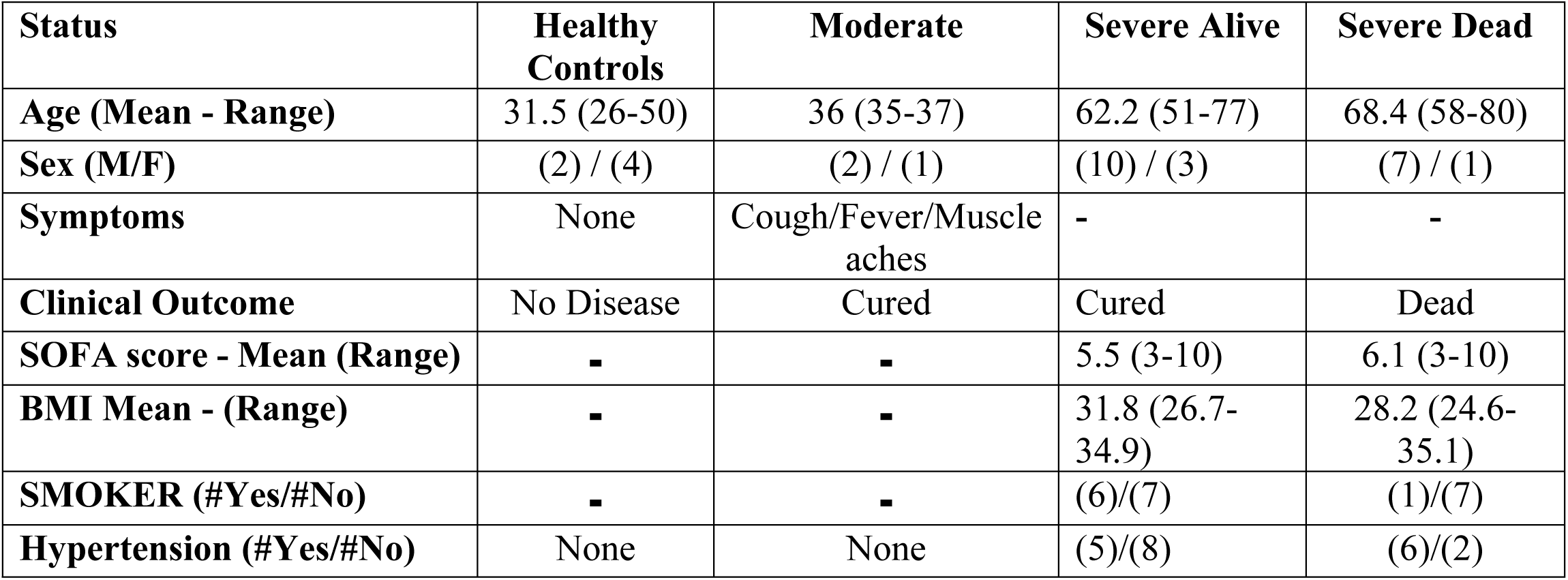
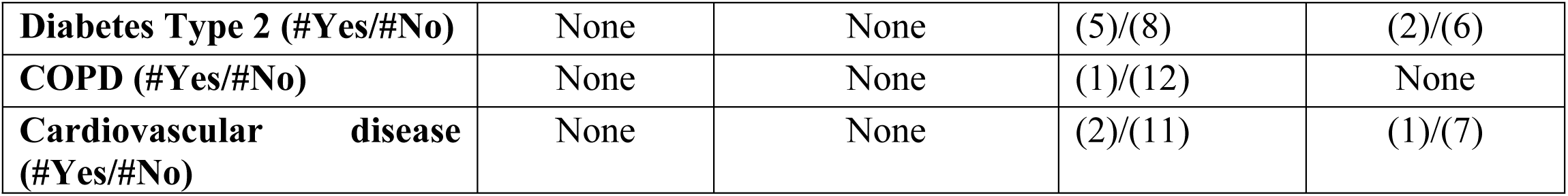
shows a detailed overview of the fundamental biological and clinical data of the individuals and patients examined in this study.

### Normalization, Feature Selection, and Clustering

For this analysis, we utilized the R package Seurat v.5.1.0[37] to perform data normalization, scaling, dimensionality reduction, clustering, differential expression analysis, and visualization. The following criteria were applied to each cell from all twenty-four patients and six healthy controls: the number of genes was restricted to between 200 and 5,000, RNA counts were capped at less than 40,000, and mitochondrial gene percentage was kept below 10%. After applying these filters, a total of 68,576 cells remained for downstream analyses. Normalization of all sample data was conducted using the LogNormalize method, followed by the selection of the top 2,000 highly variable genes for integration. For dimensionality reduction and clustering, nearest neighbor graphs (FindNeighbours) were constructed to identify cell-cell relationships. Subsequently, Louvain clustering (FindClusters) was performed with a resolution parameter of 0.8 to define distinct cellular populations. To visualize the data in two dimensions, Uniform Manifold Approximation and Projection (UMAP) was executed on the top 20 principal components. Cell clusters were annotated based on the origin of the samples, distinguishing between healthy controls, moderately ill patients, and severe COVID-19 patients. Visualization using DimPlot facilitated the exploration of clustering patterns and the identification of sample-specific clusters.

### Data Integration and Annotation

To integrate datasets from healthy controls, moderately ill patients, and severe COVID-19 patients, we utilized an anchor-based canonical correlation analysis (CCA) method. This approach aligns datasets by identifying shared sources of variation, improving comparability and addressing batch effects across different studies[38]. After integration, neighbors were recalculated based on the integrated data, and clustering was performed with a resolution of 1.2 to define integrated clusters. UMAP projection using the integrated coordinates provided a unified visualization of cell populations across conditions. Additionally, we accessed single-cell RNA sequencing (ScRNA-seq) data from the Human Primary Cell Atlas reference database using the celldex package in R. Automatic cell type annotation was conducted with the SingleR package in R, which compares the gene expression profiles of our dataset against the reference dataset from the Human Primary Cell Atlas.

### Re-clustering of Relevant Immune Cells and Differential Gene Expression

We focused on re-clustering cells identified as neutrophils, monocytes/macrophages, NK cells, and T-cells to concentrate on relevant innate immune cells and T-cells. For comparing gene expression in individual cells, the FindMarkers function was used, considering genes with an adjusted p-value < 0.05 as significantly differentially expressed. To compare differential gene expression among patient groups for each cell type, we applied Seurat’s implementation of the Wilcoxon rank-sum test, using a two-sided p-value < 0.05 and the average log(fold-change) of each differentially expressed gene.

### Data Availability

Upon acceptance and publication of this manuscript, all relevant data files, generated using GraphPad Prism version 10.2.3, will be made publicly accessible. The data can be retrieved via a dedicated FigShare repository and https://github.com. Further details and the link to the repository will be provided at the time of publication.

## Results

### Single-Cell RNA Sequencing of Bronchoalveolar Cells Reveals Differential Immune Profiles with Increasing Severity in COVID-19 Patients

The complexity and heterogeneity of immune responses to SARS-CoV-2 infection present formidable challenges in understanding the pathology of COVID-19[39, 40]. To elucidate the cellular dynamics within the lung microenvironment, we conducted a secondary data analysis of single-cell RNA sequencing (scRNA-seq) datasets derived from bronchoalveolar lavage fluid (BALF) cells. The study encompassed four distinct patient cohorts: healthy controls, individuals with moderate COVID-19 (Moderate), patients with severe COVID-19 who survived (Severe Alive - SevereA), and those with severe COVID-19 who did not survive (Severe Dead - SevereD). The use of bronchoalveolar lavage (BAL) represents a minimally invasive, established technique for sampling the lower respiratory tract, offering an in-depth view of the cellular components within the lung’s bronchioles and alveoli. This makes BALF an ideal medium for investigating lung cellular dynamics, particularly in respiratory diseases such as COVID-19.

Utilizing key cellular markers, we identified and confirmed the distribution of specific cell types: CD68 and CD14 for monocytes/macrophages, FCGR3B for neutrophils, CD56 for natural killer (NK) cells, and CD3 for T cells. These markers enabled precise characterization of the cellular landscape and facilitated differential expression analysis across the cohorts.

Our analysis delineated 29 distinct immune cell populations. The UMAP visualization (**Fig 1A&B**) revealed clear segregation of macrophages/monocytes (clusters 2, 4, 10, 11, 12, 15, 17, 19, 27), neutrophils (clusters 0, 1, 3, 5, 6, 8, 9, 14, 16), NK cells (cluster 22), and T cells (clusters 7, 13, 18, 22). Neutrophils, marked by high FCGR3B expression, were significantly elevated in severe COVID-19 cases, particularly among those who did not survive (**Fig 1C**). This elevation underscores the intense inflammatory response associated with severe disease. In moderate cases, neutrophil levels remained relatively low, akin to those in healthy controls. Conversely, macrophages/monocytes, identified by CD68 and CD14, were most abundant in healthy controls and moderate cases, with a notable reduction in severe cases (**Fig 1D**). This depletion or functional exhaustion of macrophages/monocytes in severe COVID-19 patients underscores their critical role in disease progression and immune dysregulation.

**Figure 1:**
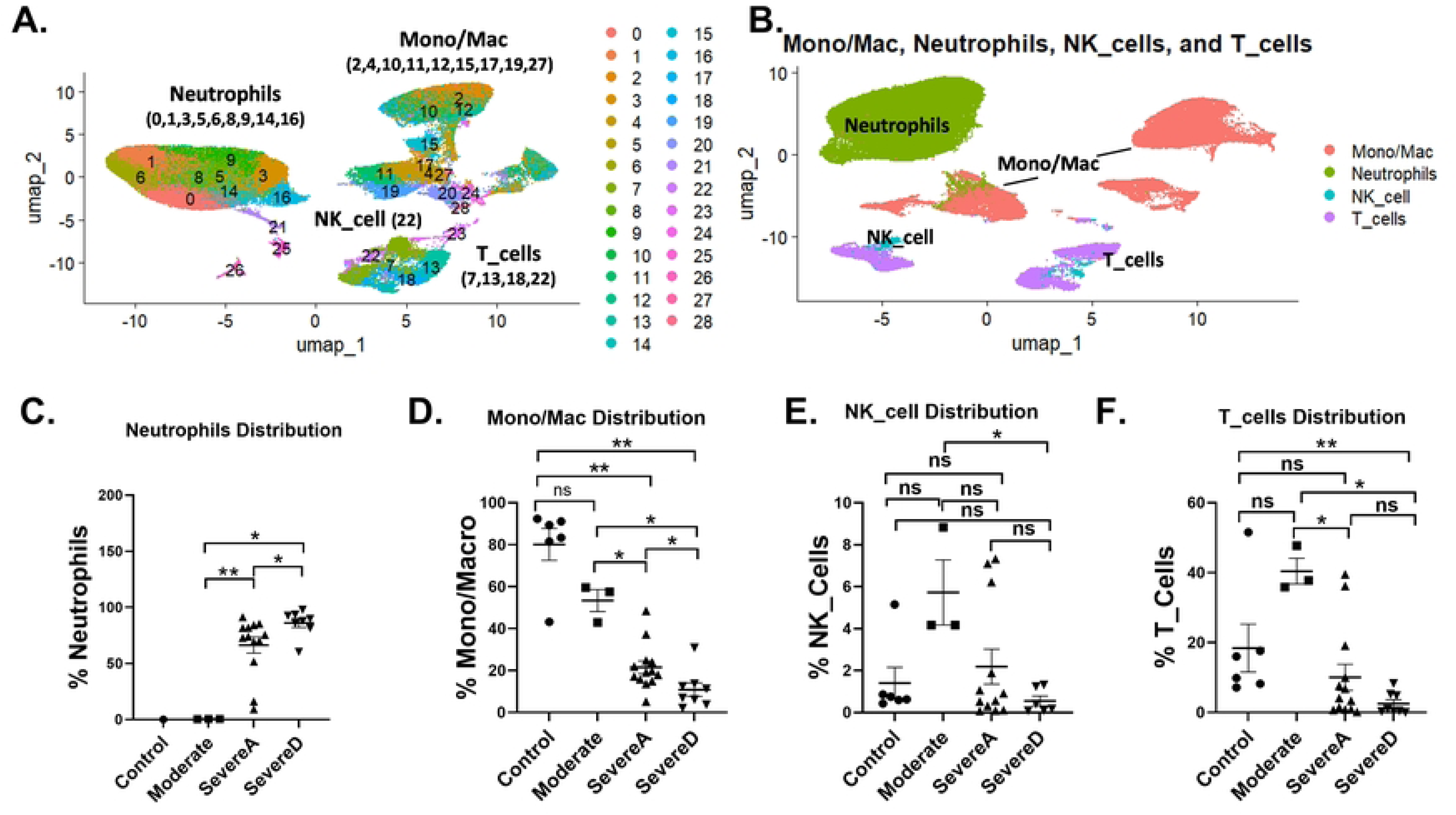
Single-Cell Atlas of Immune Subsets in Bronchoalveolar Lavage Fluid from Healthy and COVID-19 Patients. (**A**) Two-dimensional UMAP representation of the major immune cell types and associated clusters from bronchoalveolar lavage fluid among healthy donors and COVID-19 patients. (**B**) UMAP clustering distribution of four immune cell populations [Neutrophils, Monocytes-Macrophages (Mono/Mac), NK cells, and T cells]. The proportion of BALF immune cell types focusing on (**C**)Neutrophils, (**D**) Mono/Mac, (**E**) NK cells, and (**F**) T cells among Healthy Controls (n=6), Moderate COVID-19 patients (n=3), Severe Alive COVID-19 patients [SevereA] (n=13), and Severe Dead COVID-19 patients [SevereD] (n=8). Statistical analysis utilized the t-test with posttest Wilcox verification. Significance levels are indicated as follows: p < 0.001 (***), p < 0.01 (**), and p < 0.05 (*).

NK cells, characterized by CD56 expression, were generally low across all groups but exhibited a marked decrease in severe cases, suggesting a compromised innate immune response critical for early viral defense. The NK cell percentages were significantly reduced in severe cases compared to controls and moderate cases (**Fig 1E**). T cells, identified by CD3, also showed significant reductions in severe cases, reflecting an impaired adaptive immune response. The T cell depletion was particularly pronounced in patients who did not survive (**Fig 1F**), highlighting the crucial role of T cells in disease control.

The cohort analysis revealed distinct patterns in immune cell distributions (**Fig 1**). Healthy controls exhibited balanced immune cell distributions, with high levels of monocytes/macrophages and T cells, and low levels of neutrophils and NK cells. Moderate COVID-19 patients showed increased immune activation compared to controls but maintained relatively balanced proportions, with a slight rise in neutrophils and a decrease in NK cells. Severe COVID-19 patients who survived displayed drastic immune alterations, with elevated neutrophil levels and reduced monocytes/macrophages and NK cells, along with significant T cell depletion. The most profound immune dysregulation was observed in patients who did not survive, characterized by dominant neutrophil presence, minimal monocyte/macrophage and NK cell levels, and severe T cell depletion (**Fig 1C, D,E** and **F**).

### Elevated Neutrophil Activation, Trap Formation, and Survival in Bronchoalveolar Lavage Fluid Correlate with Increased Disease Severity in COVID-19

Neutrophils are crucial components of the innate immune system, playing both protective and harmful roles in inflammation and tissue damage across various diseases. They form neutrophil extracellular traps (NETs) to capture and kill pathogens, but excessive NET formation can contribute to tissue damage and exacerbate disease severity. To better understand the role of neutrophils in both protecting and damaging the respiratory system, their characteristics in bronchoalveolar lavage (BAL) fluid from healthy donors and COVID-19 patients were examined using single-cell transcriptomics data analysis.

Through UMAP data clustering of neutrophils (**Fig 2**), we identified 6 neutrophil clusters. Neutrophils from Severe Alive (SevereA) and Severe Dead (SevereD) were clustered within populations 0, 1, 2, 3, 4, 5, while populations 1, 2 and 3 showed an increase in SevereD compared to SevereA, associating these clusters with potential roles in impacting disease outcomes (**Fig 2A, B, and C**). Population 0, found in healthy controls at very low levels, showed a progressive significant increase in moderate, SevereA, and SevereD COVID-19 patients (**Fig 2B**). Populations 1, 2 and 3, which are not present in healthy controls and moderate patients, showed a significant and progressive increase in SevereA cases, with even greater elevation observed in SevereD COVID-19 patients (**Fig 2B**).

**Figure 2:**
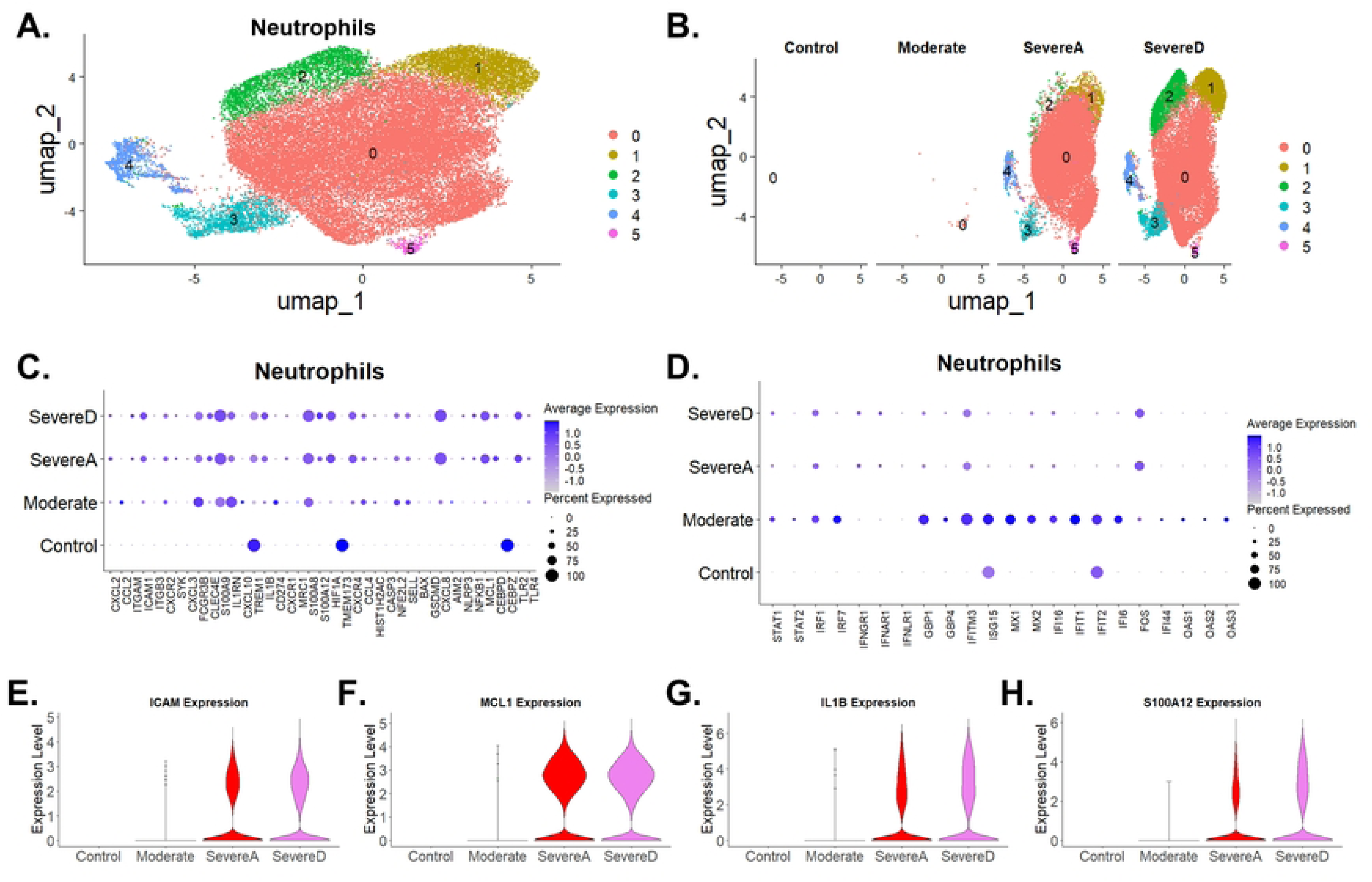
Comparative Neutrophil Cluster Trajectories in Bronchoalveolar Lavage from Healthy and COVID-19 Patients. (**A**) UMAP plot showing six neutrophil subclusters from healthy individuals and COVID-19 patients. (**B**) Proportions and distribution of the various neutrophil clusters showing differential expressions across healthy donors and three patient groups (Moderate, SevereA, and SevereD). (**C**) Dot plot illustration of gene expression profiles of neutrophils focusing on top markers for chemokines, immune and antiviral regulatory genes, inflammatory, innate immune sensing, tissue homeostasis genes, and cell death/viability genes among healthy subjects and three COVID-19 patient groups (Moderate, SevereA, and SevereD). (**D**) Expression of top marker genes involved in antiviral immune protection and interferon response regulation among healthy control subjects and three COVID-19 patient groups (Moderate, SevereA, and SevereD). (**E, F, G & H)** Violin plot illustrations of neutrophil expression of (**E**) ICAM, (**F**) MCL1, (**G**) IL1β, and (**H**) S100A12 among healthy subjects and three COVID-19 patient groups (Moderate, SevereA, and SevereD).

In healthy controls, neutrophils maintained a resting state with low expression levels of genes such as C-X-C Motif Chemokine Ligand 2 (CXCL2), C-X-C Motif Chemokine Ligand 3 (CXCL3), and S100 Calcium Binding Protein A8 (S100A8). In patients with moderate COVID-19, there was a significant upregulation of L-selectin (SELL/CD62L) and C-X-C Motif Chemokine Receptor 2 (CXCR2), indicating an activated state aimed at moderate inflammation control and pathogen clearance (**Fig 2C**).

In severe COVID-19 cases, both in survivors (SevereA) and non-survivors (SevereD), gene expression profiles shifted dramatically. Significant upregulation of chemokine ligand, MCP-1 (CCL2) and C-X-C Motif Chemokine Ligand 10 (CXCL10) increased in the moderate cases but decreased with increase disease severity. The macrophage inflammatory protein 1-beta (CCL4) increased across moderate to severe cases while C-X-C Motif Chemokine Ligand 8 (CXCL8), C-X-C Motif Chemokine Ligand 2 (CXCL2), and S100A8 increases were observed in both severe groups, pointing to an intense inflammatory response (**Fig 2C**). However, the heightened expression of Triggering Receptor Expressed on Myeloid Cells 1 (TREM1), IL-1β and Nuclear Factor Kappa B Subunit 1 (NFκB1) was more pronounced in SevereD, indicating a further intensified immune response in non-survivors. NFE2L2 expression in neutrophils was significantly higher in COVI-19 patients compared to the healthy controls (**Fig 2C**). This increased expression suggests a pivotal role of NFE2L2 in the heightened immune response seen in severe COVID-19 cases, likely as a compensatory mechanism to counteract elevated oxidative stress (**Fig 2C**). These findings indicate that NFE2L2 could serve as a biomarker for disease severity and a potential therapeutic target.

The expression of CD274 (Programmed Death-Ligand 1, PD-L1) was elevated in moderate cases but showed lower levels in SevereA and SevereD (**Fig 2C**). Upregulation of genes such as Fc Gamma Receptor IIIb (FCGR3B) and Selectin L (SELL, L-selectin) was noted in moderate cases, highlighting neutrophils’ role in adhesion and migration, facilitating their accumulation at the infection site (**Fig 2C**). However, these genes decreased in SevereA and SevereD, suggesting increased recruitment and phagocytosis in moderate cases but decreased in severe cases (**Fig 2C**).

Strikingly, our data analysis revealed reduced BCL2 Associated X, Apoptosis Regulator (BAX) and increased Myeloid Cell Leukemia 1 (MCL1), suggesting direct regulatory roles in neutrophil viability (Fig 2C). In severe COVID-19 cases, elevated levels of MCL1 indicative of enhanced neutrophil survival, with a higher expression in SevereA compared to SevereD (**Fig 2C**). Interestingly, there was a reduction in the expression of Transmembrane Protein 173 (TMEM173) in severe cases, with a more pronounced decrease in SevereD (**Fig 2C**).

Caspase 3 (CASP3), BAX, and Gasdermin D (GSDMD) did not show significant changes, suggesting that cell death pathways might not be heavily induced in these patient groups (**Fig 2C**). TREM1 expression increased in healthy controls, decreased in moderate cases, then showed a progressive increase in SevereA and SevereD, suggesting a role in modulating the immune response across disease severity (**Fig 2C**). Superoxide Dismutase (SOD) showed a progressive increase in moderate, SevereA, and SevereD, suggesting its involvement in managing oxidative stress. NLR Family Pyrin Domain Containing 3 (NLRP3) increased in SevereA, and SevereD, indicating its role in inflammasome activation and inflammation (**Fig 2C**).

Additionally, a differential expression pattern between high-density neutrophils (HDNs) and low-density neutrophils (LDNs) across patient groups was revealed. In severe cases (SevereA and SevereD), there was an increased presence of LDNs, characterized by elevated expression of C-X-C Motif Chemokine Receptor 4 (CXCR4) and CD274 (increased only in moderate and decreased in severe cases), while HDNs showed reduced activity, as indicated by lower expression levels of C-X-C Motif Chemokine Receptor 2 (CXCR2) and SELL, more so in Severe cases (**Fig 2C**).

Toll-Like Receptor 2 (TLR2) expression increased with moderate and severe COVID-19, suggesting its role in recognizing pathogens and triggering immune responses. C-Type Lectin Domain Family 4 Member E (CLEC4E) expression increased with disease severity in moderate, SevereA, and SevereD, suggesting its involvement in pathogen recognition and immune modulation (**Fig 2C**). CCAAT/Enhancer Binding Protein Zeta (CEBPZ) showed high levels in healthy controls but decreased in moderate cases and showed a tendency towards increasing in severe cases, indicating its potential role in neutrophil function and differentiation. Conversely, CCAAT/Enhancer Binding Protein Delta (CEBPD) had low levels in controls but showed a marked increase in SevereA and SevereD, suggesting its involvement in inflammation and immune response regulation (**Fig 2C**). Histone Cluster 1 H2AC (HIST1H2AC) showed increased expression in moderate cases but reduced expression with increased COVID-19 severity, suggesting its role in chromatin structure and gene regulation in the context of inflammation (**Fig 2C**).

Spleen Tyrosine Kinase (SYK) expression was very low in control cases but increased in SevereA, and SevereD, indicating its role in regulating inflammation and other pathways, which might become more relevant in severe cases (**Fig 2C**). The signal transducer and activator of transcription 1 & 2 (STAT1 and STAT2), Interferon Regulatory Factor 1 (IRF1), and Interferon Regulatory Factor 7 (IRF7), showed varied expression patterns across patient groups, highlighting their roles in antiviral responses and immune regulation (**Fig 2D**). Increased expression of Interferon Induced Transmembrane Protein 3 (IFITM3), Interferon Induced Protein with Tetratricopeptide Repeats 1 (IFIT1), Interferon Induced Protein with Tetratricopeptide Repeats 2 (IFIT2), Interferon Alpha Inducible Protein 6 &16 (IFI6 and IFI16), and guanylate binding proteins 1 & 2 (GBP1 and GBP2) was observed in moderate cases, correlating with heightened antiviral immune activation (**Fig 2D**). In addition, other interferon induced genes such as Oligoadenylate Synthetase 1 (OAS1), Oligoadenylate Synthetase 2 (OAS2), Oligoadenylate Synthetase 3 (OAS3), Myxovirus resistance 1 and 2 (MX1 and MX2) showed increased expression in moderate cases, emphasizing their roles in antiviral defense mechanisms (**Fig 2D**).

Next, we concentrated on evaluating the significant upregulation of genes vital to immune response and inflammation in neutrophils. Intercellular Adhesion Molecule 1 (ICAM1) increased notably in severe COVID-19 patients who survived, with an even greater rise in those who died, highlighting its role in leukocyte adhesion and migration during severe inflammation (**Fig 2E**). Similarly, Myeloid Cell Leukemia 1 (MCL1) showed a progressive increase from severe alive to severe dead patients, indicating its involvement in cell survival and apoptosis regulation during severe infection (**Fig 2F**). Interleukin 1 Beta (IL1β) also exhibited a substantial rise from severe alive to severe dead patients, underscoring its role in mediating fever and systemic inflammation (**Fig 2G**). S100 Calcium Binding Protein A12 (S100A12) followed a similar trend, emphasizing its contribution to the inflammatory response and disease progression (**Fig 2H**). The consistent increase in ICAM1, MCL1, IL1β, and S100A12 expression in neutrophils from severe alive to severe dead patients underscores the critical role of neutrophils in COVID-19 severity. This pattern reflects a heightened and potentially dysregulated immune response in neutrophils, contributing to poorer prognosis in severe cases.

### Heightened Inflammatory Macrophage Phenotypes with Progressive Disease Severity in COVID-19 Patients from Single Cell Analysis of Bronchoalveolar Lavage

Macrophages are pivotal in immune responses, recognizing and destroying pathogens, aiding tissue repair, and modulating inflammation. Disruptions in these processes can lead to poor pathogen clearance and excessive inflammation, causing tissue damage. We conducted a single-cell transcriptomics analysis of bronchoalveolar lavage (BAL) samples from healthy individuals and COVID-19 patients to investigate how monocyte-macrophage functions vary with disease severity. By further sub-setting monocytes and macrophages (Mono_Mac), we identified eight macrophage subclusters, highlighting the diversity among patient groups (**Fig 3A**).

**Figure 3:**
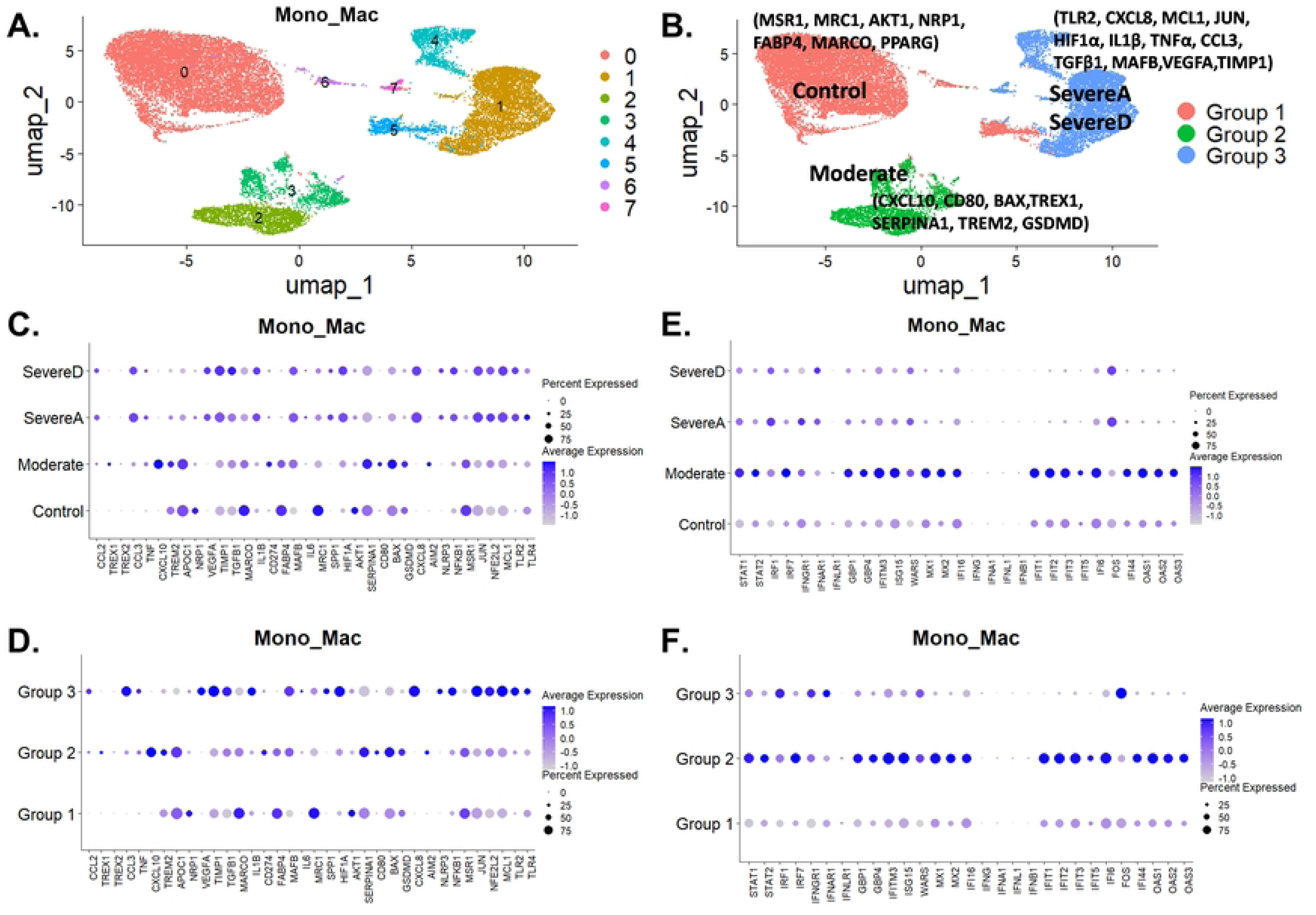
Diverse Single Cell Landscape of Macrophage Variations between Healthy Individuals and COVID-19 Patients. (**A**) Clustering of macrophages into 8 groups from healthy and COVID-19 infected patients. (**B**) Subgrouping revealing three major clusters with distinct gene profiles linking distributions to Healthy Controls (n=6) -Group1, Moderate COVID-19 patients (n=3) -Group 2, Severe Alive COVID-19 patients (SevereA) (n=13) -Group 3, and Severe Dead COVID-19 patients (SevereD) (n=8) -Group 3. (**C & D**) Expression of top marker genes involved in chemokines, immune and antiviral regulation, inflammation, innate immune sensing, tissue homeostasis, and cell death/viability among healthy individuals and the three COVID-19 patient groups (Moderate, SevereA, and SevereD) shown in dot plots. (**E & F**) Dot plot illustration of the expression profiles of top marker genes involved in antiviral immune protection and interferon response regulation among healthy individuals and three COVID-19 patient groups (Moderate, SevereA, and SevereD), with gene profile distributions to the three groupings in (B).

Through UMAP subgrouping of macrophage populations, we identified three primary clusters: Group 1, Group 2, and Group 3, associating these clusters with patient groups, including healthy controls (**Fig 3B**). Group 1 mainly comprised healthy controls, Group 2 consisted primarily of patients with mild COVID-19, and Group 3 predominantly included patients with severe COVID-19 (**Fig 3B**).

In healthy controls (Group 1), monocyte-macrophages showed baseline expression of key inflammatory markers like Tumor Necrosis Factor alpha (TNFα) and Interleukin 1 beta (IL1β), with higher levels of anti-inflammatory markers such as Peroxisome Proliferator-Activated Receptor Gamma (PPAR-γ), Mannose Receptor C Type 1 (MRC1), and Macrophage Receptor with Collagenous Structure (MARCO), suggesting a regulatory and tissue repair phenotype (**Fig 3C**). Additionally, genes like Cluster of Differentiation 274 (CD274), Fatty Acid Binding Protein 4 (FABP4), and Macrophage Scavenger Receptor 1 (MSR1) had higher expression levels, reinforcing an anti-inflammatory and tissue repair function (**Fig 3C**).

With increasing COVID-19 severity, the expression of inflammatory markers intensified. In moderate cases (Group 2), genes such as C-X-C Motif Chemokine Ligand 8 (CXCL8), Apolipoprotein C1 (APOC1), Bcl-2-associated X protein (BAX), Serpin Family A Member 1 (SERPINA1), Triggering Receptor Expressed on Myeloid Cells 2 (TREM2), and Gasdermin D (GSDMD) were highly expressed, linking this group to inflammation and cell stress responses (**Fig 3C**). Three Prime Repair Exonuclease 1 (TREX1) expression also showed an increase in moderate cases, suggesting a potential capacity for macrophages to degrade neutrophil NETs. Elevated chemokines like C-C Motif Chemokine Ligand 2 (CCL2) and C-C Motif Chemokine Ligand 3 (CCL3) suggested increased recruitment of more immune cells to the infection site. Moreover, genes like Transforming Growth Factor Beta 1 (TGFβ1), Tissue Inhibitor of Metalloproteinases 1 (TIMP1), and Vascular Endothelial Growth Factor A (VEGFA) had increased expression in severe cases (SevereA and SevereD), indicating a shift towards an anti-inflammatory state (**Fig 3C**). Increased GSDMD expression in moderate cases suggested potential cell stress and pyroptosis, with decreases in severe cases suggesting reduced pyroptosis. TREM2 upregulation in moderate cases indicated an increased capacity for macrophages to clear damage associated molecular patterns (DAMPs) and apoptotic cells via enhanced phagocytosis/efferocytosis. However, TREM2 expression decreased in severe cases (SevereA and SevereD), suggesting decreased clearance of DAMPs/apoptotic cells in these groups (**Fig 3C**).

Group 3 predominantly expressed genes like Toll-Like Receptor 2 (TLR2), CXCL8, Myeloid Cell Leukemia Sequence 1 (MCL1), Jun Proto-Oncogene, AP-1 Transcription Factor Subunit (JUN), Hypoxia Inducible Factor 1 Subunit Alpha (HIF1α), IL1β, TNFα, CCL3, TGFβ1, and MAF BZIP Transcription Factor B (MAFB), reflecting a highly inflammatory and hypoxia-adapted phenotype (**Fig 3C**). In severe cases, especially in SevereA and SevereD groups, there was significant upregulation of pro-inflammatory markers such as IL1β, TNFα, and Interleukin 6 (IL6). Genes involved in hypoxia and metabolic adaptation, such as HIF1α, were notably elevated in severe cases, with more pronounced expression in SevereD. Decreased SERPINA1 levels in severe cases indicated impaired anti-protease activity, particularly in SevereD (**Fig 3C**). Decreased phagocytosis in severe COVID-19 was evident by the downregulation of MSR1, Neuropilin 1 (NRP1), and MRC1, with more significant decreases in SevereD (**Fig 3C**). Elevated JUN expression in severe cases suggested increased cellular stress responses and potential promotion of inflammatory gene expression. NFE2L2 expression was elevated in macrophages from patients with severe disease compared to controls (**Fig 3C**). This suggests that NFE2L2 plays a crucial role in macrophage function and the inflammatory response during severe COVID-19. The gene’s involvement in modulating antioxidant responses and inflammation underscores its potential as a target for therapeutic interventions aimed at mitigating the cytokine storm and tissue damage in severe cases.

Additionally, increased expression of inflammasome-related markers like Absent In Melanoma 2 (AIM2), NLR Family Pyrin Domain Containing 3 (NLRP3), and GSDMD in monocyte-macrophages suggested a higher incidence of pyroptosis during severe SARS-CoV-2 compared to moderate infection (**Fig 3C**). Activation of the NLRP3 inflammasome pathway was linked to enhanced TLRs-NF-κB axis activation. Elevated levels of TLRs such as TLR2 and Toll-Like Receptor 4 (TLR4), alongside increased expression of the Nuclear Factor Kappa B (NF-κB) transcription factor, were observed more so in severe compared to moderate COVID-19 patients (**Fig 3C**). NF-κB activation downstream of TLR signaling pathways drives the expression of pro-inflammatory cytokines like IL-1β, TNFα, and IL-6, especially prominent in severe SARS-CoV-2 infections.

Our analysis also identified distinct monocyte-macrophage phenotypes, specifically M1 (pro-inflammatory) and M2 (anti-inflammatory), across patient groups. In severe cases, M1 monocyte-macrophages dominated, indicated by elevated expression of IL1β, TNFα, and IL6, with higher levels in SevereA (**Fig 3C**). Conversely, M2 monocyte-macrophages, characterized by higher expression of MSR1, MRC1, and PPAR-γ, were more prevalent in control and moderate cases. PPAR-γ promotes monocyte-macrophages towards an M2 phenotype with increased expression of anti-inflammatory genes like Interleukin 10 (IL-10) and TGF-β (**Fig 3C**).

Distinct gene expression patterns were also observed across different patient groups. In the control group, key genes such as MSR1, MRC1, PPAR-γ, CD274, FABP4, and MARCO were upregulated, indicating a tissue repair and anti-inflammatory profile. Moderate cases showed increased expression of CXCL8, APOC1, TREM2, TGFβ1, TIMP1, and VEGFA, indicating an intermediate inflammatory response (Fig 3D). Severe cases exhibited very high levels of TLR2, HIF1α, IL1β, TNFα, CCL3, TLR4, and NLRP3, reflecting a hypoxia-adapted and highly inflammatory phenotype (Fig 2C). Additionally, the increased expression of genes related to hypoxia (HIF1α), pyroptosis (NLRP3, IL1 β), and the NF-κB pathway (TLR2, TLR4, NF-κB) in Group 3 from mostly SevereA and SevereD patients underscores the severe inflammatory response and cellular stress in these patients (**Fig 3C**). The gene expression profiling of inflammatory genes, chemokines, inflammasomes, and immune sensing genes in both healthy controls and COVID patients (**Fig 3C**) showed a close correlation in expression profiles. These profiles align with the three macrophage clusters’ unique gene profiles, which link distinct macrophage subpopulations to either healthy donors, moderate COVID-19, or severe COVID-19 (**Fig 3B**).

Next, we evaluated gene expression profiles for top genes modulating Antiviral Response, Interferon Signaling, and Interferon-Stimulated Genes (ISGs) (**Fig 3E**). We found that genes such as Signal Transducer and Activator of Transcription 1 (STAT1) and STAT2, Interferon Regulatory Factor 1 (IRF1) and IRF7 play crucial roles in antiviral responses and immune regulation. Interferon Gamma Receptor 1 (IFNGR1), Interferon Alpha and Beta Receptor Subunit 1 (IFNAR1), Interleukin 1 Receptor (INFLR1), and ISGs like Interferon-Induced Transmembrane Protein 3 (IFITM3), ISG15 Ubiquitin-Like Modifier (ISG15), Tryptophanyl-tRNA Synthetase (WARS), Myxovirus Resistance Proteins 1 and 2 (MX1 and MX2), Interferon Gamma Inducible Protein 16 (IFI16), and Oligoadenylate Synthetases 1, 2, and 3 (OAS1, OAS2, OAS3), which are pivotal in interferon signaling and antiviral defense, increased markedly in moderate COVID-19 patients. However, a dramatic decrease in their expression was observed in severe COVID-19 cases, suggesting an impaired immune response and potential immune exhaustion. Conversely, FOS, which was low in healthy and moderate patients, showed a dramatic increase in severe COVID-19 cases, indicating heightened inflammation and ineffective interferon antiviral responses (**Fig 3F**). Strikingly, the gene expression profiling of the top modulating Antiviral Response, Interferon Signaling, and Interferon-Stimulated Genes (ISGs) in both healthy controls and COVID patients (**Fig 3E**) showed a close correlation in expression profiles. These profiles align with the three macrophage clusters’ unique gene profiles, which link distinct macrophage subpopulations to either healthy donors, moderate COVID-19, or severe COVID-19 (**Fig 3B**).

### Infection with SARS-CoV2 is associated with Differential Gene Expression in NK Cells that can impact COVID-19 severity

Natural Killer (NK) cells are crucial elements of the immune system, essential for controlling viral infections. These cells can detect and eliminate virus-infected cells without prior sensitization by releasing cytotoxic granules containing perforin and granzymes, which trigger apoptosis in infected cells. Additionally, NK cells produce cytokines such as interferon-gamma (IFNγ), which possess potent antiviral properties and help modulate the adaptive immune response, enhancing the body’s ability to combat infections. Our single-cell analysis of bronchoalveolar lavage (BAL) samples from healthy individuals and COVID-19 patients reveals distinct clustering patterns correlating with disease severity. These clusters highlight the heterogeneity in NK cell response among different severity groups, indicating differential expression profiles among Control, Moderate, SevereA, and SevereD patient groups (**Fig 4A**).

**Figure 4:**
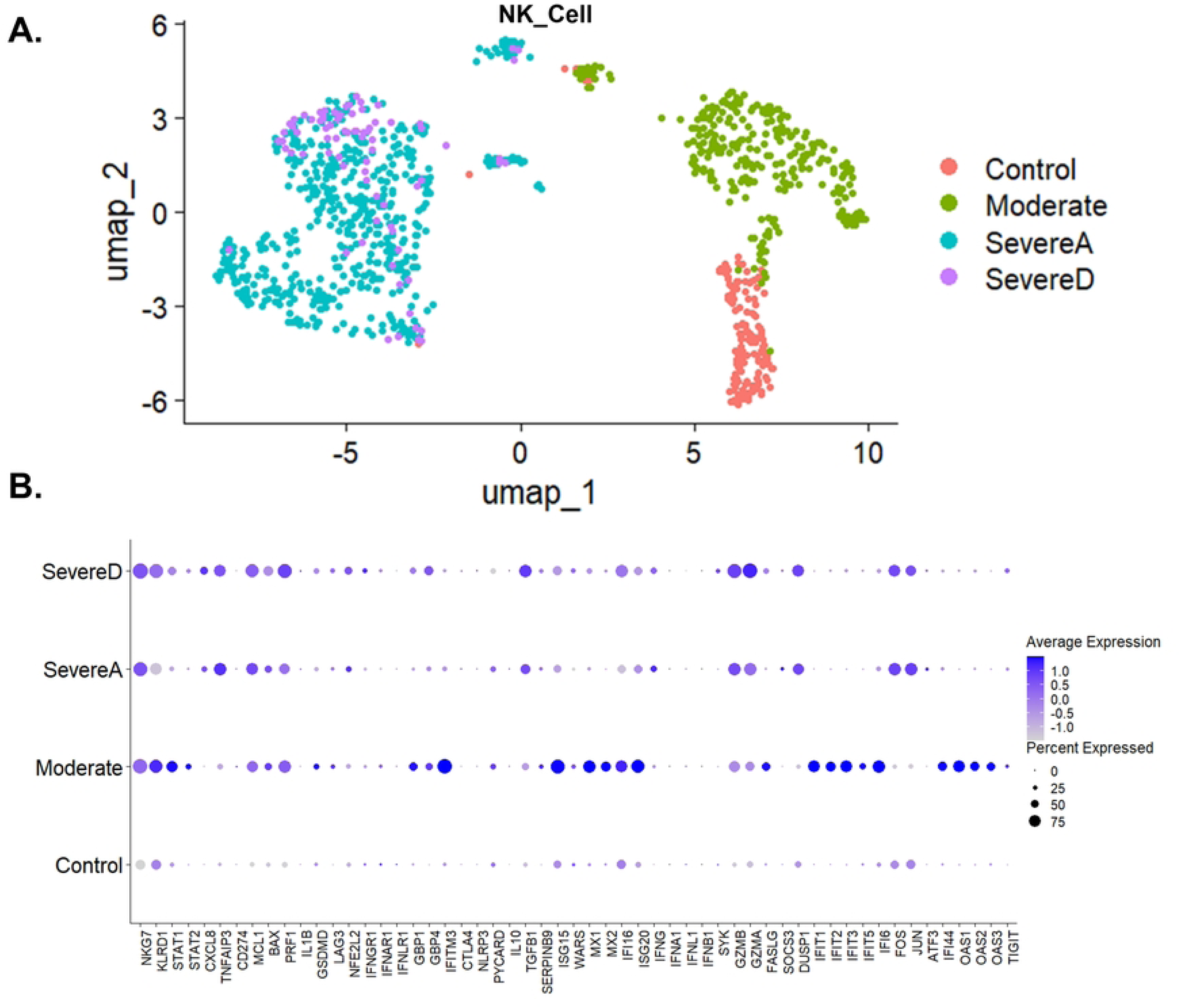
Gene Expression Profiling of NK Cell in Bronchoalveolar Lavage Fluid from Healthy Individuals and COVID-19 Patients. (**A**) UMAP distribution of NK cells among healthy control individuals and the three COVID-19 patient groups (Moderate, SevereA, and SevereD). (**B**) Dot plot representation of top gene expression profiles of NK cells from healthy individuals and the three groups of COVID-19 patients (Moderate, SevereA, and SevereD).

In severe COVID-19 cases (both SevereA and SevereD), NK cells exhibit significantly higher expression levels of IFNγ, while interferon-stimulated gene 15 (ISG15), and Myxovirus resistance protein 1 (MX1) were highly expressed in moderate cases (**Fig 4B**). IFNγ, a critical cytokine in antiviral responses and immune regulation, is markedly upregulated in SevereA, while its expression in SevereD is elevated but not as pronounced, suggesting potential dysregulation or exhaustion of NK cells in fatal cases (**Fig 4B**). ISG15 and MX1, both interferon-stimulated genes essential for antiviral defense, show significant upregulation in moderate cases, indicating an intensified antiviral response, which, although crucial for controlling the virus, can lead to tissue damage and inflammation if not properly regulated (**Fig 4B**).

Chemokine (C-X-C motif) ligand 8 (CXCL8, also known as IL-8) is notably higher in SevereD compared to SevereA, Moderate, and Control groups (**Fig 4B**). This chemokine plays a role in neutrophil recruitment to the lungs, contributing to the inflammatory response observed in severe COVID-19 cases. Interferon-induced transmembrane protein 3 (IFITM3) and Guanylate binding proteins (GBP1 and GBP4), which play roles in antiviral defense and immune regulation, are upregulated in moderate cases (**Fig 4B**).

Negative regulators of inflammation, such as TNFα induced protein 3 (TNFAIP3), which acts as a negative regulator of NF-κB signaling and inflammation, are upregulated in SevereA compared to Moderate and Control groups, suggesting a feedback mechanism attempting to limit excessive inflammation in severe cases (**Fig 4B**). Similarly, stress response and apoptosis-related genes FOS, JUN, and activating transcription factor 3 (ATF3) show increased expression in severe cases, highlighting the role of NK cells in modulating cell death pathways in response to viral infection (**Fig 4B**). Nuclear factor, erythroid 2-like 2 (NFE2L2) expression is markedly increased in NK cells from patients with severe COVID-19, indicating its role in influencing NK cell activity and antiviral responses, likely reflecting an attempt to mitigate oxidative stress and inflammation (**Fig 4B**). Dual specificity phosphatase 1 (DUSP1) and suppressor of cytokine signaling 1 (SOCS3), involved in the negative regulation of cytokine signaling and immune responses, exhibit differential expression with upregulation in Severe cases, indicating attempts by NK cells to control excessive inflammation (**Fig 4B**).

Perforin 1 (PRF1) and granzyme A/B (GZMA/B), critical for NK cell cytotoxic functions, exhibit differential expression with upregulation in SevereA and SevereD, indicating enhanced cytotoxic activity in severe cases (**Fig 4B**). This highlights the increased potential for NK cells to induce apoptosis in virus-infected cells in severe COVID-19. Killer cell lectin-like receptor subfamily D member 1 (KLRD1) and natural killer cell granule protein 7 (NKG7), involved in NK cell activation and degranulation, are upregulated in moderate and severe cases, suggesting enhanced activation and degranulation activity, essential for the cytotoxic function of NK cells during SARS-CoV-2 infection (**Fig 4B**).

Inhibitory receptors, such as lymphocyte activating 3 (LAG3) and T cell immunoreceptor with Ig and ITIM domains (TIGIT), show differential expression with higher levels in moderate compared to severe cases, indicating NK cell ability to induce increased inhibitory signaling in moderate COVID-19 but not in fatal cases. SOCS3 shows upregulation in SevereA compared to SevereD, playing a critical role in controlling the cytokine signaling cascade and preventing excessive inflammatory responses. The decreased expression of SOCS3 in the non survivors suggest a failed attempt of NK cells to mitigate the inflammatory damage associated with severe COVID-19 (**Fig 4B**).

Finally, in individuals with moderate SARS-CoV-2 infection, 2’-5’-Oligoadenylate Synthetase 1 (OAS1), OAS2, OAS3, and T cell Immunoreceptor with Ig and ITIM domains (TIGIT) were elevated, but these levels significantly decreased with increased COVID-19 severity (**Fig 4B**). OAS1, OAS2, and OAS3 are crucial in the antiviral response by degrading viral RNA, while TIGIT regulates NK cell activity to prevent excessive inflammation. The dramatic decrease in these genes with increased severity suggests a compromised antiviral defense and a dysregulated NK cell response.

### Differential T-cell Gene Expression Correlates with Severity and T Cell Dysfunctions in COVID-19

T cells are a crucial component of the adaptive immune system, playing a pivotal role in the antiviral response. These cells identify and destroy virus-infected cells, coordinate immune responses, and provide long-term immunity. T cell functions include cytotoxic activities mediated by cytotoxic T lymphocytes (CTLs), helper functions carried out by helper T cells (Th), and regulatory functions mediated by regulatory T cells (Tregs). The effectiveness of T cells in antiviral responses is governed by a complex interplay of gene expressions that control their activation, function, exhaustion, and survival.

In the context of COVID-19, we analyzed single-cell transcriptomics data from bronchoalveolar lavage fluid (BALF) across different patient groups: healthy controls (Control), moderate infection (Moderate), SevereAlive (SevereA), and SevereDead (SevereD). We focused on key genes involved in T cell functions, evaluating their expression levels and implications for disease severity, with sub clustering based on classical T cell markers.

**Fig 5A** shows the total cell distribution across different patient groups. In healthy controls, we observed a balanced distribution of T cell populations. In moderate cases, there was a notable shift towards increased cytotoxic T lymphocyte (CTL) populations. Severe cases, both SevereA and SevereD, displayed disrupted cell distributions, indicating significant immune dysregulation.

**Figure 5:**
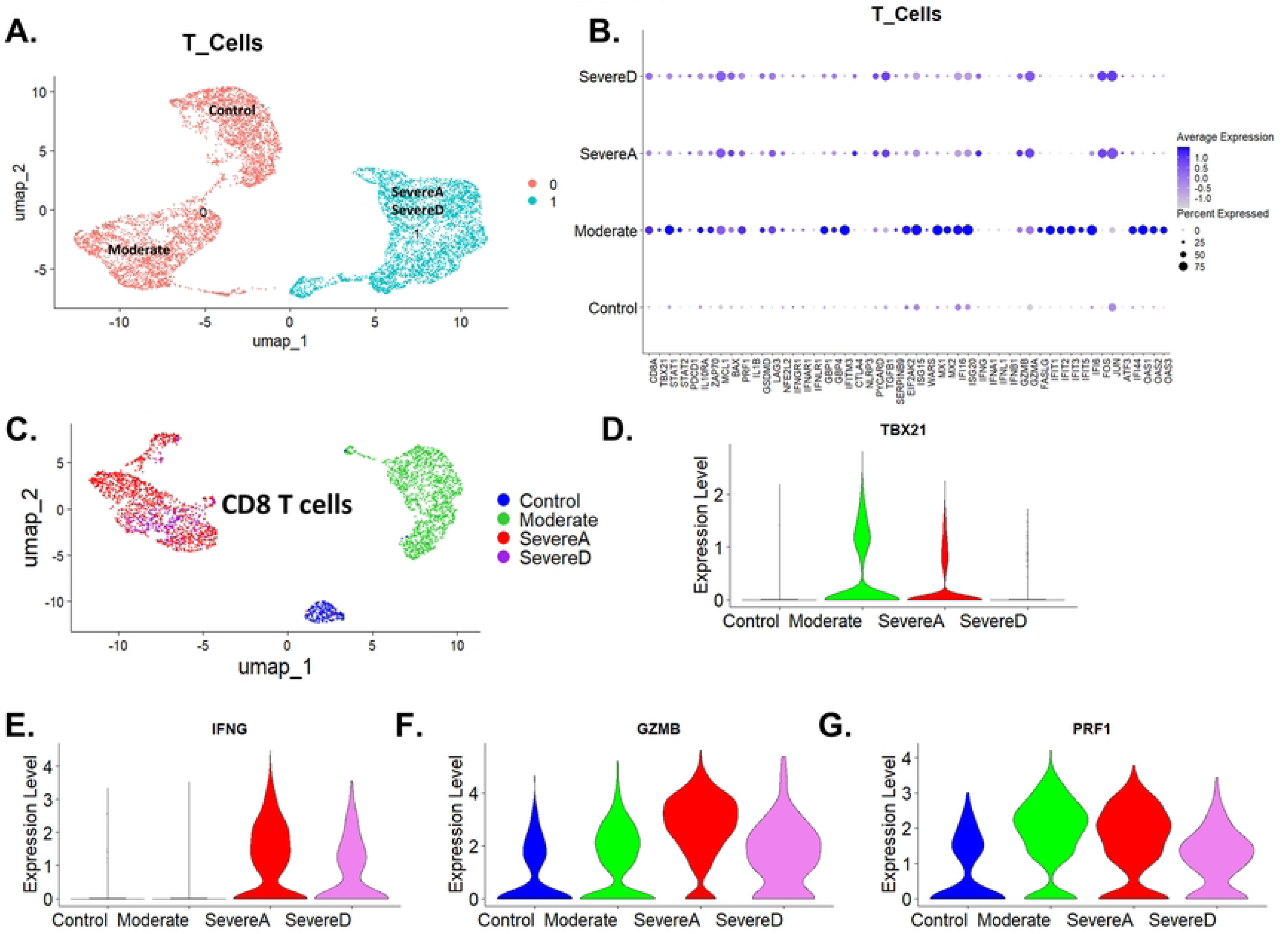
Gene Expression Alterations in T Cells from Bronchoalveolar Lavage Fluid of Healthy and COVID-19 Patients. (**A**) UMAP plot showing T cell distribution among healthy individuals (Control) and COVID-19 patients (Moderate, SevereA, and SevereD). (**B**) Dot plot representation of top gene expression profiles of T cells from healthy individuals and the three groups of COVID-19 patients (Moderate, SevereA, and SevereD). (**C**) UMAP of CD8 T cells showing the distribution of cells in distinct groups among healthy subjects and COVID-19 patients (Moderate, SevereA, and SevereD). (**D, E, F, & G**) Violin plot illustrations of the four top genes associated with direct antiviral functions of CD8 cells, including (**D**) TBX21, (**E**) IFNγ, (**F**) Granzyme B (GZMB), and (**G**) Perforin (PRF1).

**Fig 5B** shows the gene distribution in total T cells across Control, Moderate, SevereA, and SevereD COVID-19 patients. The expression of CD8 alpha chain (CD8A) is minimal in the Control group, suggesting baseline expression in T cells from healthy individuals. In the Moderate group, there is a notable increase in CD8A expression, indicating an active cytotoxic T cell response. CD8A expression decreases in SevereA but remains elevated relative to Control. In SevereD, CD8A expression is further diminished, indicating a significant reduction in cytotoxic T cell activity in the most severe cases.

Perforin 1 (PRF1), granzyme A (GZMA), and granzyme B (GZMB) are elevated in the Moderate group, highlighting increased cytotoxic activity crucial for the destruction of virus-infected cells (**Fig 5B**). In SevereA and SevereD, the expression of these genes decreases, reflecting a reduced cytotoxic response. T-box transcription factor 21 (TBX21), also known as T-bet, is significantly increased in the Moderate group, promoting the development of effective CTLs and influencing Th1 differentiation. Its expression decreases in SevereA and SevereD, indicating impaired CTL function in severe disease.

Signal transducer and activator of transcription 1 (STAT1) and signal transducer and activator of transcription 2 (STAT2) show higher expression in the Moderate group, suggesting a regulated antiviral response. Their expression is reduced in SevereA and SevereD, indicating impaired antiviral signaling. Interferon gamma (IFNG) levels are notably higher in severe cases, reflecting an intense inflammatory response. Elevated IFNG levels in SevereA and SevereD correlate with increased expression of its receptor IFNγ R1 on macrophages, enhancing their activation.

Markers of immune regulation and exhaustion show significant changes across the patient groups (**Fig 5B**). Programmed cell death protein 1 (PDCD1 or PD-1) expression is elevated in SevereA and SevereD, indicating T cell exhaustion. Interleukin 10 receptor subunit alpha (IL10RA) and zeta chain of T cell receptor-associated protein kinase 70 (ZAP70) show increased expression in the Moderate group, suggesting enhanced cytokine signaling and immune regulation. In SevereA and SevereD, the expression of these genes is reduced, indicating compromised immune regulation. Cytotoxic T-lymphocyte-associated protein 4 (CTLA4) is elevated in severe cases, indicating attempts at immune regulation. Lymphocyte activation gene 3 (LAG3) is upregulated in moderate and severe cases, reflecting immune activation regulation.

Genes involved in apoptosis and cell survival also show differential expression (**Fig 5B**). Myeloid cell leukemia 1 (MCL1), an anti-apoptotic gene, is upregulated in severe cases, potentially reflecting survival signals from the microenvironment, whereas BCL2 associated X (BAX) is upregulated, indicating ongoing cellular apoptosis. Transforming growth factor beta 1 (TGFB1) is elevated in severe cases, indicating attempts to regulate the hyperinflammatory environment.

Inflammatory markers and genes involved in the inflammasome pathway show significant upregulation in moderate and severe cases (**Fig 5B**). Interleukin 1 beta (IL1β) and tumor necrosis factor alpha (TNFα) are substantially upregulated in the Moderate group, indicating a strong inflammatory response. Their expression decreases in SevereA and SevereD, reflecting a diminished inflammatory response in severe cases. Gasdermin D (GSDMD) follows a similar expression pattern, being elevated in Moderate cases and reduced in SevereA and SevereD. NLR family pyrin domain containing 3 (NLRP3) and PYD and CARD domain containing (PYCARD or ASC) are significantly higher in severe groups, suggesting robust inflammasome activation contributing to the severe inflammatory response.

Interferon-stimulated genes and antiviral response genes exhibit distinct patterns (**Fig 5B**). Interferon-induced transmembrane protein 3 (IFITM3) and guanylate-binding proteins 1 and 4 (GBP1 and GBP4) show minimal expression in controls but are significantly upregulated in the moderate infection, indicating a strong antiviral response. Their expression decreases in SevereA and SevereD, reflecting a weakened antiviral response in severe disease. Interferon-stimulated genes such as ISG15 ubiquitin-like modifier (ISG15), tryptophanyl-tRNA synthetase (WARS), MX dynamin-like GTPase 1 and 2 (MX1 and MX2), and interferon gamma-inducible protein 16 (IFI16) are upregulated in moderate cases, indicating robust antiviral responses, but their expression decreases in SevereA and SevereD. Similar patterns are observed for interferon-stimulated gene 20 (ISG20), interferon alpha 1 (IFNA1), interferon lambda 1 (IFNL1), interferon beta 1 (IFNB1), interferon-induced proteins with tetratricopeptide repeats 1, 2, 3, 5, and 6 (IFIT1, IFIT2, IFIT3, IFIT5, and IFIT6), Fos proto-oncogene (FOS), Jun proto-oncogene (JUN), activating transcription factor 3 (ATF3), and interferon-induced protein 44 (IFI44). Oligoadenylate synthetase 1, 2, and 3 (OAS1, OAS2, and OAS3) show minimal expression in Control but are significantly upregulated in Moderate cases, indicating a strong antiviral response. Their expression decreases in SevereA and SevereD, reflecting a weakened antiviral response.

Given the direct antiviral role of CD8 T cells, we then went further to subset and characterize these cells in more detail. **Fig 5C** shows the UMAP distribution of CD8 T cells according to different patient groups. The analysis reveals distinct clustering patterns, reflecting functional heterogeneity and varying responses to SARS-CoV-2 infection. We further analyzed the expression of key genes involved in CD8 T cell functions: TBX21 (T-bet), IFNG, GZMB, and PRF1.

The violin plots in **Figs 5D, E, F, and G** reveal critical information on CD8 T cell functions across different patient groups compared to controls. TBX21 (T-bet) expression is significantly elevated in Moderate cases, promoting effective CTL responses. In SevereA and SevereD, reduced T-bet expression indicates impaired CTL differentiation and function. IFNG is elevated in severe cases, underscoring the intense inflammatory response and its role in immune-mediated pathology. GZMB expression is increased in Moderate cases, correlating with enhanced cytotoxic activity. Decreased expression in SevereA and SevereD reflects diminished CTL function. PRF1 follows a similar pattern, being elevated in Moderate cases, supporting effective cytotoxic responses, but decreasing in SevereA and SevereD, indicating compromised CTL-mediated cytotoxicity.

## Discussion

Despite the availability of multiple vaccines and antiviral treatments, COVID-19 continues to pose formidable and recurring threats to human health globally. Understanding the evolving pathology is crucial for better disease management. Our extensive single-cell RNA sequencing (scRNA-seq) analysis of secondary bronchoalveolar lavage (BAL) single-cell transcriptomics data from COVID-19 patients and healthy controls reveals novel insights into the immune landscape of the lung during SARS-CoV-2 infection. Our findings highlight critical immune dysregulation correlating with disease severity, offering groundbreaking perspectives on COVID-19 pathogenesis and potential therapeutic targets. By elucidating the complex interplay among neutrophils, macrophages, NK cells, and T cells in the context of COVID-19, this study significantly advances the understanding of cellular dynamics within the lung microenvironment during this devastating disease.

The severity of COVID-19 cases, as indicated by clinical parameters, demonstrated higher mean ages and a greater prevalence of comorbidities such as hypertension, diabetes, and cardiovascular disease compared to healthy controls and moderate cases. Additionally, body mass index (BMI) values were elevated in severe cases, indicating a potential link between higher body mass and disease severity. Smoking status and Sequential Organ Failure Assessment (SOFA) scores, available only for severe cases, highlighted significant organ dysfunction, with higher SOFA scores correlating with worse outcomes, particularly in non-survivors. These clinical observations are supported by previous studies that have identified similar trends, underscoring the importance of these factors in the progression and severity of COVID-19[41–46].

Our analysis of neutrophils indicates their substantial elevation in severe COVID-19 cases, particularly in non-survivors. This observation aligns with previous studies suggesting that neutrophil extracellular traps (NETs) contribute to lung damage and thrombosis in COVID-19 patients[47]. Neutrophils from severe COVID-19 cases exhibit a hyperinflammatory profile, with upregulated expression of IL1β, CXCL10, and S100A8, corroborating the findings of cytokine storm syndromes reported in severe COVID-19[48]. The elevated expression of TREM1 and NFκB1 in non-survivors suggests an exaggerated immune response that could exacerbate lung injury, as TREM1 amplifies inflammatory signaling[49, 50]. This enhanced inflammatory response in severe and fatal cases of COVID-19 indicates that neutrophil activity is not merely a marker of disease severity but may actively contribute to the pathogenesis of severe disease. Moreover, the differential expression patterns observed in moderate versus severe cases suggest that while neutrophils play a protective role in moderate COVID-19 by controlling inflammation and aiding in pathogen clearance, their excessive activation in severe cases might results in excessive tissue damage and worse clinical outcomes.

Furthermore, we observed differential neutrophil subpopulations, with an increased presence of low-density neutrophils (LDNs) in severe cases, particularly non-survivors, adding another layer of complexity to our findings. LDNs have been associated with chronic inflammation and immunosuppression in various diseases[51]. The heightened expression of CXCR4 and CD274 (PD-L1) in LDNs indicates a potential mechanism for immune evasion and persistent inflammation, adding a new dimension to the understanding of neutrophil heterogeneity in COVID-19. These findings suggest that targeting LDNs and their signaling pathways could be a promising therapeutic strategy to mitigate severe COVID-19 outcomes. The presence of these immunosuppressive neutrophil subsets may contribute to the impaired antiviral responses observed in severe COVID-19, further complicating the immune landscape. By delineating the specific roles and states of neutrophil subsets, this study offers a detailed map of neutrophil dynamics, providing a foundation for targeted therapies that could selectively modulate harmful neutrophil functions without compromising their protective roles.

Transitioning to the role of macrophages, these cells play a pivotal part in orchestrating immune responses and maintaining tissue homeostasis. Our data reveal a shift towards a pro-inflammatory M1 phenotype in severe COVID-19 cases, with elevated expression of TNFα, IL1β, CCL2, and CCL3, which are markers of classical activation and inflammation. This shift is more pronounced in the severe group compared to the moderate group, suggesting that persistent M1 activation contributes to severe lung pathology[52]. The upregulation of genes involved in hypoxia (HIF1α) and metabolic adaptation (NFE2L2) in macrophages from severe COVID-19 cases indicates a metabolic reprogramming that supports their inflammatory function. Additionally, the increased expression of pyroptosis-related genes (GSDMB, AIM2, NLRP3) during SARS-CoV-2 infection highlights cell death pathways contributing to lung damage and inflammation. This reprogramming enhances the macrophages’ ability to sustain prolonged inflammation, contributing to the severe pathology observed in these patients.[53]. This is supported by studies showing that pyroptosis is a key mechanism of immune-mediated tissue injury in COVID-19[54, 55] . The reduced expression of phagocytic receptors (MSR1, MRC1) in severe cases, especially non-survivors, suggests impaired macrophage clearance of apoptotic cells and debris, potentially exacerbating inflammation, and tissue damage. This aligns with the concept of “cytokine storm” where dysregulated macrophage activity leads to overwhelming inflammation[10, 56, 57].

Building on these insights, our data on macrophages offer crucial insights into the progressive immune dysregulation seen in COVID-19. The transition from M2 to M1 macrophages indicates a failure to resolve inflammation, which is critical for healing and recovery. This finding is corroborated by similar shifts observed in other inflammatory diseases and underscores the potential for therapies aimed at rebalancing macrophage phenotypes. The metabolic reprogramming of macrophages towards glycolysis in severe cases aligns with the hypoxic conditions of inflamed tissues, suggesting that interventions targeting metabolic pathways could modulate macrophage function and ameliorate disease severity. Furthermore, the distinct expression profiles of macrophages in moderate versus severe COVID-19 provide a potential biomarker for disease progression and a target for therapeutic intervention to prevent the escalation of inflammation and tissue damage[58].

Transitioning to NK cells, these cell types are critical for antiviral immunity and have shown significant dysregulation in severe COVID-19 cases. The marked decrease in NK cells, coupled with the upregulation of activation markers (IFNγ, ISG15), indicates a paradoxical state of activation and exhaustion. This is consistent with studies reporting NK cell dysfunction and depletion in severe COVID-19[48]. The reduced NK cell numbers and function likely impair early viral clearance, contributing to uncontrolled viral replication and severe disease. In moderate cases, NK cells exhibit a balanced profile, maintaining effective antiviral responses without progressing to exhaustion, highlighting the importance of NK cell function in controlling disease progression[24, 59]. The detailed profiling of NK cell subsets reveals how SARS-CoV-2 manipulates the innate immune response, providing insights into potential therapeutic targets aimed at restoring NK cell function and enhancing early antiviral defense.

Similarly, T cells, particularly CD8+ cytotoxic T cells, showed significant reductions in severe cases, indicating another layer of immune dysregulation. This reduction, coupled with increased expression of exhaustion markers (PDCD1, CTLA4), suggests that T cells in severe COVID-19 are overactivated and subsequently exhausted, impairing their ability to control the virus. This finding is supported by previous research showing T cell exhaustion in chronic viral infections and severe COVID-19[60–62] . The profound T cell dysregulation observed highlights the dual challenge of managing inflammation and restoring effective antiviral immunity. Therapeutic strategies that focus on reversing T cell exhaustion and restoring functional T cell responses are critical for improving outcomes in severe COVID-19 patients[63, 64]. Furthermore, the role of NFE2L2 (Nuclear Factor, Erythroid 2-Like 2), commonly known as NRF2, adds another dimension to understanding the oxidative stress response in severe COVID-19 cases. NRF2 is a transcription factor that regulates the expression of antioxidant proteins that protect against oxidative damage triggered by injury and inflammation. Our data show increased expression of NRF2-related genes in severe cases, suggesting an adaptive response to the heightened oxidative stress in these patients. This aligns with recent studies highlighting the potential protective role of NRF2 activation in mitigating the severe inflammatory and oxidative damage seen in COVID-19[65, 66]. The modulation of NRF2 pathways could represent a therapeutic strategy to enhance antioxidant defenses and reduce tissue damage in severe COVID-19.

The immune dysregulation observed highlights potential therapeutic targets for COVID-19, with multiple facets requiring attention. Modulating neutrophil activity and preventing NET formation could mitigate lung damage and thrombosis[47]. Therapies targeting TREM1 and NFκB signaling may reduce excessive inflammation and improve outcomes in severe cases. Enhancing macrophage phagocytic function and preventing pyroptosis could also be beneficial[53]. Drugs that stabilize macrophage metabolism and promote M2 polarization might help restore immune balance and reduce lung injury. For NK and T cells, strategies to prevent exhaustion and restore function are critical. Immune checkpoint inhibitors targeting PD-1/PD-L1 may rejuvenate exhausted T cells, enhancing antiviral responses. Additionally, therapies that boost NK cell numbers and function could improve early viral clearance and prevent severe disease progression[48].

Our novel findings underscore the importance of the interplay between different immune cell types in COVID-19 pathogenesis. The heightened inflammatory response driven by neutrophils likely exacerbates macrophage dysregulation, leading to a vicious cycle of inflammation and tissue damage. The observed reduction in NK cells and T cells further impairs the immune system’s ability to control viral replication and resolve inflammation[58] . This intricate interaction among immune cells highlights the need for a multi-faceted therapeutic approach that addresses the dysregulation of multiple cell types to effectively manage severe COVID-19.

In conclusion, our analysis of secondary BAL single-cell transcriptomics data from COVID-19 patients provides a detailed view of the immune landscape in the lung, revealing critical dysregulation correlating with disease severity. The novel insights into neutrophil heterogeneity, macrophage phenotypes, and NK/T cell dysfunction underscore the complexity of the immune response in COVID-19. These findings not only enhance the understanding of COVID-19 pathogenesis but also identify potential targets with diagnostic, prognostic, and potential therapeutic intervention. Integrating clinical data, including SOFA scores, BMI, and comorbidities, underscores the relevance of these findings to disease severity and patient outcomes. Continued research integrating single-cell transcriptomics with clinical data will be essential to develop effective treatments and improve patient outcomes in COVID-19 and other pro-inflammatory respiratory viral infections. By providing a detailed map of immune cell dynamics and identifying key dysregulated pathways, this study offers a foundation for targeted therapies that could selectively modulate harmful immune functions without compromising protective roles. This approach holds promise for improving clinical outcomes and managing severe COVID-19 and other respiratory viral infections more effectively.

## References

1. Ritchie H ME, Rodés-Guirao L, Appel C, Giattino C, Ortiz-Ospina E, et al. Coronavirus Pandemic (COVID-19). Our World in Data. 2024;Retrived July 14th 2024.

2. Malireddi RKS, Sharma BR, Kanneganti TD. Innate Immunity in Protection and Pathogenesis During Coronavirus Infections and COVID-19. Annu Rev Immunol. 2024;42(1):615–45. Epub 2024/06/28. doi: 10.1146/annurev-immunol-083122-043545. PubMed PMID: 38941608.

3. Cheong JG, Ravishankar A, Sharma S, Parkhurst CN, Grassmann SA, Wingert CK, et al. Epigenetic memory of coronavirus infection in innate immune cells and their progenitors. Cell. 2023;186(18):3882–902 e24. Epub 20230818. doi: 10.1016/j.cell.2023.07.019. PubMed PMID: 37597510; PubMed Central PMCID: PMCPMC10638861.

4. Newton AH, Cardani A, Braciale TJ. The host immune response in respiratory virus infection: balancing virus clearance and immunopathology. Semin Immunopathol. 2016;38(4):471–82. Epub 2016/03/12. doi: 10.1007/s00281-016-0558-0. PubMed PMID: 26965109; PubMed Central PMCID: PMCPMC4896975.

5. See H, Wark P. Innate immune response to viral infection of the lungs. Paediatr Respir Rev. 2008;9(4):243–50. Epub 2008/11/26. doi: 10.1016/j.prrv.2008.04.001. PubMed PMID: 19026365; PubMed Central PMCID: PMCPMC7172072.

6. Rosales C. Neutrophil: A Cell with Many Roles in Inflammation or Several Cell Types? Front Physiol. 2018;9:113. Epub 2018/03/09. doi: 10.3389/fphys.2018.00113. PubMed PMID: 29515456; PubMed Central PMCID: PMCPMC5826082.

7. Chan L, Karimi N, Morovati S, Alizadeh K, Kakish JE, Vanderkamp S, et al. The Roles of Neutrophils in Cytokine Storms. Viruses. 2021;13(11). Epub 2021/11/28. doi: 10.3390/v13112318. PubMed PMID: 34835125; PubMed Central PMCID: PMCPMC8624379.

8. Kaplan MJ, Radic M. Neutrophil extracellular traps: double-edged swords of innate immunity. J Immunol. 2012;189(6):2689–95. Epub 2012/09/08. doi: 10.4049/jimmunol.1201719. PubMed PMID: 22956760; PubMed Central PMCID: PMCPMC3439169.

9. Borges L, Pithon-Curi TC, Curi R, Hatanaka E. COVID-19 and Neutrophils: The Relationship between Hyperinflammation and Neutrophil Extracellular Traps. Mediators Inflamm. 2020;2020:8829674. Epub 2020/12/22. doi: 10.1155/2020/8829674. PubMed PMID: 33343232; PubMed Central PMCID: PMCPMC7732408 publication of this paper.

10. Zanza C, Romenskaya T, Manetti AC, Franceschi F, La Russa R, Bertozzi G, et al. Cytokine Storm in COVID-19: Immunopathogenesis and Therapy. Medicina (Kaunas). 2022;58(2). Epub 20220118. doi: 10.3390/medicina58020144. PubMed PMID: 35208467; PubMed Central PMCID: PMCPMC8876409.

11. Chen S, Saeed A, Liu Q, Jiang Q, Xu H, Xiao GG, et al. Macrophages in immunoregulation and therapeutics. Signal Transduct Target Ther. 2023;8(1):207. Epub 20230522. doi: 10.1038/s41392-023-01452-1. PubMed PMID: 37211559; PubMed Central PMCID: PMCPMC10200802.

12. Murray PJ. Macrophage Polarization. Annu Rev Physiol. 2017;79:541–66. Epub 20161021. doi: 10.1146/annurev-physiol-022516-034339. PubMed PMID: 27813830.

13. Roszer T. Understanding the Mysterious M2 Macrophage through Activation Markers and Effector Mechanisms. Mediators Inflamm. 2015;2015:816460. Epub 20150518. doi: 10.1155/2015/816460. PubMed PMID: 26089604; PubMed Central PMCID: PMCPMC4452191.

14. Arish M, Qian W, Narasimhan H, Sun J. COVID-19 immunopathology: From acute diseases to chronic sequelae. J Med Virol. 2023;95(1):e28122. Epub 20220913. doi: 10.1002/jmv.28122. PubMed PMID: 36056655; PubMed Central PMCID: PMCPMC9537925.

15. Sun L, Liu S, Chen ZJ. SnapShot: pathways of antiviral innate immunity. Cell. 2010;140(3):436–e2. doi: 10.1016/j.cell.2010.01.041. PubMed PMID: 20144765; PubMed Central PMCID: PMCPMC3586550.

16. Lang PA, Lang KS, Xu HC, Grusdat M, Parish IA, Recher M, et al. Natural killer cell activation enhances immune pathology and promotes chronic infection by limiting CD8+ T-cell immunity. Proc Natl Acad Sci U S A. 2012;109(4):1210–5. Epub 20111213. doi: 10.1073/pnas.1118834109. PubMed PMID: 22167808; PubMed Central PMCID: PMCPMC3268324.

17. Cook KD, Waggoner SN, Whitmire JK. NK cells and their ability to modulate T cells during virus infections. Crit Rev Immunol. 2014;34(5):359–88. doi: 10.1615/critrevimmunol.2014010604. PubMed PMID: 25404045; PubMed Central PMCID: PMCPMC4266186.

18. Schuster IS, Coudert JD, Andoniou CE, Degli-Esposti MA. “Natural Regulators“: NK Cells as Modulators of T Cell Immunity. Front Immunol. 2016;7:235. Epub 20160614. doi: 10.3389/fimmu.2016.00235. PubMed PMID: 27379097; PubMed Central PMCID: PMCPMC4905977.

19. Lee AJ, Ashkar AA. The Dual Nature of Type I and Type II Interferons. Front Immunol. 2018;9:2061. Epub 20180911. doi: 10.3389/fimmu.2018.02061. PubMed PMID: 30254639; PubMed Central PMCID: PMCPMC6141705.

20. Ruetsch C, Brglez V, Cremoni M, Zorzi K, Fernandez C, Boyer-Suavet S, et al. Functional Exhaustion of Type I and II Interferons Production in Severe COVID-19 Patients. Front Med (Lausanne). 2020;7:603961. Epub 20210127. doi: 10.3389/fmed.2020.603961. PubMed PMID: 33585507; PubMed Central PMCID: PMCPMC7873370.

21. Lee HR, Kim MH, Lee JS, Liang C, Jung JU. Viral interferon regulatory factors. J Interferon Cytokine Res. 2009;29(9):621–7. doi: 10.1089/jir.2009.0067. PubMed PMID: 19715458; PubMed Central PMCID: PMCPMC2956608.

22. van der Ploeg EK, Carreras Mascaro A, Huylebroeck D, Hendriks RW, Stadhouders R. Group 2 Innate Lymphoid Cells in Human Respiratory Disorders. J Innate Immun. 2020;12(1):47–62. Epub 20190206. doi: 10.1159/000496212. PubMed PMID: 30726833; PubMed Central PMCID: PMCPMC6959098.

23. Xiong L, Nutt SL, Seillet C. Innate lymphoid cells: More than just immune cells. Front Immunol. 2022;13:1033904. Epub 20221026. doi: 10.3389/fimmu.2022.1033904. PubMed PMID: 36389661; PubMed Central PMCID: PMCPMC9643152.

24. Bjorkstrom NK, Strunz B, Ljunggren HG. Natural killer cells in antiviral immunity. Nat Rev Immunol. 2022;22(2):112–23. Epub 20210611. doi: 10.1038/s41577-021-00558-3. PubMed PMID: 34117484; PubMed Central PMCID: PMCPMC8194386.

25. Moretta A, Marcenaro E, Parolini S, Ferlazzo G, Moretta L. NK cells at the interface between innate and adaptive immunity. Cell Death Differ. 2008;15(2):226–33. Epub 20070601. doi: 10.1038/sj.cdd.4402170. PubMed PMID: 17541426.

26. Okeke EB, Uzonna JE. The Pivotal Role of Regulatory T Cells in the Regulation of Innate Immune Cells. Front Immunol. 2019;10:680. Epub 20190409. doi: 10.3389/fimmu.2019.00680. PubMed PMID: 31024539; PubMed Central PMCID: PMCPMC6465517.

27. Guan T, Zhou X, Zhou W, Lin H. Regulatory T cell and macrophage crosstalk in acute lung injury: future perspectives. Cell Death Discov. 2023;9(1):9. Epub 20230116. doi: 10.1038/s41420-023-01310-7. PubMed PMID: 36646692; PubMed Central PMCID: PMCPMC9841501.

28. Chevrier S, Zurbuchen Y, Cervia C, Adamo S, Raeber ME, de Souza N, et al. A distinct innate immune signature marks progression from mild to severe COVID-19. Cell Rep Med. 2021;2(1):100166. Epub 20201226. doi: 10.1016/j.xcrm.2020.100166. PubMed PMID: 33521697; PubMed Central PMCID: PMCPMC7817872.

29. Morrissey SM, Geller AE, Hu X, Tieri D, Ding C, Klaes CK, et al. A specific low-density neutrophil population correlates with hypercoagulation and disease severity in hospitalized COVID-19 patients. JCI Insight. 2021;6(9). Epub 20210510. doi: 10.1172/jci.insight.148435. PubMed PMID: 33986193; PubMed Central PMCID: PMCPMC8262329.

30. Lv J, Wang Z, Qu Y, Zhu H, Zhu Q, Tong W, et al. Distinct uptake, amplification, and release of SARS-CoV-2 by M1 and M2 alveolar macrophages. Cell Discov. 2021;7(1):24. Epub 20210413. doi: 10.1038/s41421-021-00258-1. PubMed PMID: 33850112; PubMed Central PMCID: PMCPMC8043100.

31. Kosyreva A, Dzhalilova D, Lokhonina A, Vishnyakova P, Fatkhudinov T. The Role of Macrophages in the Pathogenesis of SARS-CoV-2-Associated Acute Respiratory Distress Syndrome. Front Immunol. 2021;12:682871. Epub 20210510. doi: 10.3389/fimmu.2021.682871. PubMed PMID: 34040616; PubMed Central PMCID: PMCPMC8141811.

32. Gao M, Liu Y, Guo M, Wang Q, Wang Y, Fan J, et al. Regulatory CD4(+) and CD8(+) T cells are negatively correlated with CD4(+) /CD8(+) T cell ratios in patients acutely infected with SARS-CoV-2. J Leukoc Biol. 2021;109(1):91–7. Epub 20200915. doi: 10.1002/JLB.5COVA0720-421RR. PubMed PMID: 32930458; PubMed Central PMCID: PMCPMC10016821.

33. Moss P. The T cell immune response against SARS-CoV-2. Nat Immunol. 2022;23(2):186–93. Epub 20220201. doi: 10.1038/s41590-021-01122-w. PubMed PMID: 35105982.

34. Lim J, Park C, Kim M, Kim H, Kim J, Lee DS. Advances in single-cell omics and multiomics for high-resolution molecular profiling. Exp Mol Med. 2024;56(3):515–26. Epub 20240305. doi: 10.1038/s12276-024-01186-2. PubMed PMID: 38443594; PubMed Central PMCID: PMCPMC10984936.

35. Gondal MN, Shah SUR, Chinnaiyan AM, Cieslik M. A Systematic Overview of Single-Cell Transcriptomics Databases, their Use cases, and Limitations. ArXiv. 2024. Epub 20240415. PubMed PMID: 38699169; PubMed Central PMCID: PMCPMC11065044.

36. Liao M, Liu Y, Yuan J, Wen Y, Xu G, Zhao J, et al. Single-cell landscape of bronchoalveolar immune cells in patients with COVID-19. Nat Med. 2020;26(6):842–4. Epub 20200512. doi: 10.1038/s41591-020-0901-9. PubMed PMID: 32398875.

37. Hao Y, Stuart T, Kowalski MH, Choudhary S, Hoffman P, Hartman A, et al. Dictionary learning for integrative, multimodal and scalable single-cell analysis. Nat Biotechnol. 2024;42(2):293–304. Epub 20230525. doi: 10.1038/s41587-023-01767-y. PubMed PMID: 37231261; PubMed Central PMCID: PMCPMC10928517.

38. Stuart T, Butler A, Hoffman P, Hafemeister C, Papalexi E, Mauck WM, 3rd, et al. Comprehensive Integration of Single-Cell Data. Cell. 2019;177(7):1888–902 e21. Epub 20190606. doi: 10.1016/j.cell.2019.05.031. PubMed PMID: 31178118; PubMed Central PMCID: PMCPMC6687398.

39. Harne R, Williams B, Abdelal HFM, Baldwin SL, Coler RN. SARS-CoV-2 infection and immune responses. AIMS Microbiol. 2023;9(2):245–76. Epub 20230329. doi: 10.3934/microbiol.2023015. PubMed PMID: 37091818; PubMed Central PMCID: PMCPMC10113164.

40. da Silva Torres MK, Bichara CDA, de Almeida M, Vallinoto MC, Queiroz MAF, Vallinoto I, et al. The Complexity of SARS-CoV-2 Infection and the COVID-19 Pandemic. Front Microbiol. 2022;13:789882. Epub 20220210. doi: 10.3389/fmicb.2022.789882. PubMed PMID: 35222327; PubMed Central PMCID: PMCPMC8870622.

41. Huang C, Wang Y, Li X, Ren L, Zhao J, Hu Y, et al. Clinical features of patients infected with 2019 novel coronavirus in Wuhan, China. Lancet. 2020;395(10223):497–506. Epub 20200124. doi: 10.1016/S0140-6736(20)30183-5. PubMed PMID: 31986264; PubMed Central PMCID: PMCPMC7159299.

42. Fayed M, Patel N, Angappan S, Nowak K, Vasconcelos Torres F, Penning DH, Chhina AK. Sequential Organ Failure Assessment (SOFA) Score and Mortality Prediction in Patients With Severe Respiratory Distress Secondary to COVID-19. Cureus. 2022;14(7):e26911. Epub 20220716. doi: 10.7759/cureus.26911. PubMed PMID: 35865183; PubMed Central PMCID: PMCPMC9290429.

43. Gao M, Piernas C, Astbury NM, Hippisley-Cox J, O’Rahilly S, Aveyard P, Jebb SA. Associations between body-mass index and COVID-19 severity in 6.9 million people in England: a prospective, community-based, cohort study. Lancet Diabetes Endocrinol. 2021;9(6):350–9. Epub 20210428. doi: 10.1016/S2213-8587(21)00089-9. PubMed PMID: 33932335; PubMed Central PMCID: PMCPMC8081400.

44. Gallus S, Scala M, Possenti I, Jarach CM, Clancy L, Fernandez E, et al. The role of smoking in COVID-19 progression: a comprehensive meta-analysis. Eur Respir Rev. 2023;32(167). Epub 20230308. doi: 10.1183/16000617.0191-2022. PubMed PMID: 36889786; PubMed Central PMCID: PMCPMC10032583.

45. Wu Z, McGoogan JM. Characteristics of and Important Lessons From the Coronavirus Disease 2019 (COVID-19) Outbreak in China: Summary of a Report of 72 314 Cases From the Chinese Center for Disease Control and Prevention. JAMA. 2020;323(13):1239–42. doi: 10.1001/jama.2020.2648. PubMed PMID: 32091533.

46. Floyd JS, Walker RL, Kuntz JL, Shortreed SM, Fortmann SP, Bayliss EA, et al. Association Between Diabetes Severity and Risks of COVID-19 Infection and Outcomes. J Gen Intern Med. 2023;38(6):1484–92. Epub 20230216. doi: 10.1007/s11606-023-08076-9. PubMed PMID: 36795328; PubMed Central PMCID: PMCPMC9933797.

47. Barnes BJ, Adrover JM, Baxter-Stoltzfus A, Borczuk A, Cools-Lartigue J, Crawford JM, et al. Targeting potential drivers of COVID-19: Neutrophil extracellular traps. J Exp Med. 2020;217(6). doi: 10.1084/jem.20200652. PubMed PMID: 32302401; PubMed Central PMCID: PMCPMC7161085.

48. Mathew D, Giles JR, Baxter AE, Oldridge DA, Greenplate AR, Wu JE, et al. Deep immune profiling of COVID-19 patients reveals distinct immunotypes with therapeutic implications. Science. 2020;369(6508). Epub 20200715. doi: 10.1126/science.abc8511. PubMed PMID: 32669297; PubMed Central PMCID: PMCPMC7402624.

49. Bouchon A, Facchetti F, Weigand MA, Colonna M. TREM-1 amplifies inflammation and is a crucial mediator of septic shock. Nature. 2001;410(6832):1103-7. doi: 10.1038/35074114. PubMed PMID: 11323674.

50. Wu X, Zeng H, Xu C, Chen H, Fan L, Zhou H, et al. TREM1 Regulates Neuroinflammatory Injury by Modulate Proinflammatory Subtype Transition of Microglia and Formation of Neutrophil Extracellular Traps via Interaction With SYK in Experimental Subarachnoid Hemorrhage. Front Immunol. 2021;12:766178. Epub 20211013. doi: 10.3389/fimmu.2021.766178. PubMed PMID: 34721438; PubMed Central PMCID: PMCPMC8548669.

51. Silvin A, Chapuis N, Dunsmore G, Goubet AG, Dubuisson A, Derosa L, et al. Elevated Calprotectin and Abnormal Myeloid Cell Subsets Discriminate Severe from Mild COVID-19. Cell. 2020;182(6):1401–18 e18. Epub 20200805. doi: 10.1016/j.cell.2020.08.002. PubMed PMID: 32810439; PubMed Central PMCID: PMCPMC7405878.

52. Hu T, Pang N, Li Z, Xu D, Jing J, Li F, et al. The Activation of M1 Macrophages is Associated with the JNK-m6A-p38 Axis in Chronic Obstructive Pulmonary Disease. Int J Chron Obstruct Pulmon Dis. 2023;18:2195–206. Epub 20231006. doi: 10.2147/COPD.S420471. PubMed PMID: 37822331; PubMed Central PMCID: PMCPMC10564081.

53. Arunachalam PS, Wimmers F, Mok CKP, Perera R, Scott M, Hagan T, et al. Systems biological assessment of immunity to mild versus severe COVID-19 infection in humans. Science. 2020;369(6508):1210-20. Epub 20200811. doi: 10.1126/science.abc6261. PubMed PMID: 32788292; PubMed Central PMCID: PMCPMC7665312.

54. Wang M, Chang W, Zhang L, Zhang Y. Pyroptotic cell death in SARS-CoV-2 infection: revealing its roles during the immunopathogenesis of COVID-19. Int J Biol Sci. 2022;18(15):5827–48. Epub 20220921. doi: 10.7150/ijbs.77561. PubMed PMID: 36263178; PubMed Central PMCID: PMCPMC9576507.

55. Bittner ZA, Schrader M, George SE, Amann R. Pyroptosis and Its Role in SARS-CoV-2 Infection. Cells. 2022;11(10). Epub 20220523. doi: 10.3390/cells11101717. PubMed PMID: 35626754; PubMed Central PMCID: PMCPMC9140030.

56. Lange A, Lange J, Jaskula E. Cytokine Overproduction and Immune System Dysregulation in alloHSCT and COVID-19 Patients. Front Immunol. 2021;12:658896. Epub 20210602. doi: 10.3389/fimmu.2021.658896. PubMed PMID: 34149697; PubMed Central PMCID: PMCPMC8206782.

57. Fajgenbaum DC, June CH. Cytokine Storm. N Engl J Med. 2020;383(23):2255–73. doi: 10.1056/NEJMra2026131. PubMed PMID: 33264547; PubMed Central PMCID: PMCPMC7727315.

58. Lucas C, Wong P, Klein J, Castro TBR, Silva J, Sundaram M, et al. Longitudinal analyses reveal immunological misfiring in severe COVID-19. Nature. 2020;584(7821):463-9. Epub 20200727. doi: 10.1038/s41586-020-2588-y. PubMed PMID: 32717743; PubMed Central PMCID: PMCPMC7477538.

59. Zuo W, Zhao X. Natural killer cells play an important role in virus infection control: Antiviral mechanism, subset expansion and clinical application. Clin Immunol. 2021;227:108727. Epub 20210420. doi: 10.1016/j.clim.2021.108727. PubMed PMID: 33887436; PubMed Central PMCID: PMCPMC8055501.

60. Zheng HY, Zhang M, Yang CX, Zhang N, Wang XC, Yang XP, et al. Elevated exhaustion levels and reduced functional diversity of T cells in peripheral blood may predict severe progression in COVID-19 patients. Cell Mol Immunol. 2020;17(5):541–3. Epub 20200317. doi: 10.1038/s41423-020-0401-3. PubMed PMID: 32203186; PubMed Central PMCID: PMCPMC7091621.

61. Rha MS, Shin EC. Activation or exhaustion of CD8(+) T cells in patients with COVID-19. Cell Mol Immunol. 2021;18(10):2325–33. Epub 20210819. doi: 10.1038/s41423-021-00750-4. PubMed PMID: 34413488; PubMed Central PMCID: PMCPMC8374113.

62. Kusnadi A, Ramirez-Suastegui C, Fajardo V, Chee SJ, Meckiff BJ, Simon H, et al. Severely ill COVID-19 patients display impaired exhaustion features in SARS-CoV-2-reactive CD8(+) T cells. Sci Immunol. 2021;6(55). doi: 10.1126/sciimmunol.abe4782. PubMed PMID: 33478949; PubMed Central PMCID: PMCPMC8101257.

63. Diao B, Wang C, Tan Y, Chen X, Liu Y, Ning L, et al. Reduction and Functional Exhaustion of T Cells in Patients With Coronavirus Disease 2019 (COVID-19). Front Immunol. 2020;11:827. Epub 20200501. doi: 10.3389/fimmu.2020.00827. PubMed PMID: 32425950; PubMed Central PMCID: PMCPMC7205903.

64. Schub D, Klemis V, Schneitler S, Mihm J, Lepper PM, Wilkens H, et al. High levels of SARS-CoV-2-specific T cells with restricted functionality in severe courses of COVID-19. JCI Insight. 2020;5(20). Epub 20201015. doi: 10.1172/jci.insight.142167. PubMed PMID: 32937615; PubMed Central PMCID: PMCPMC7605520.

65. Calabrese EJ, Kozumbo WJ, Kapoor R, Dhawan G, Lara PC, Giordano J. Nrf2 activation putatively mediates clinical benefits of low-dose radiotherapy in COVID-19 pneumonia and acute respiratory distress syndrome (ARDS): Novel mechanistic considerations. Radiother Oncol. 2021;160:125–31. Epub 20210428. doi: 10.1016/j.radonc.2021.04.015. PubMed PMID: 33932453; PubMed Central PMCID: PMCPMC8080499.

66. Cuadrado A, Pajares M, Benito C, Jimenez-Villegas J, Escoll M, Fernandez-Gines R, et al. Can Activation of NRF2 Be a Strategy against COVID-19? Trends Pharmacol Sci. 2020;41(9):598–610. Epub 20200714. doi: 10.1016/j.tips.2020.07.003. PubMed PMID: 32711925; PubMed Central PMCID: PMCPMC7359808.

